# Moxifloxacin-mediated killing of *Mycobacterium tuberculosis* involves respiratory downshift, reductive stress, and ROS accumulation

**DOI:** 10.1101/2022.04.04.486929

**Authors:** Somnath Shee, Samsher Singh, Ashutosh Tripathi, Chandrani Thakur, Anand Kumar T, Mayashree Das, Vikas Yadav, Sakshi Kohli, Raju S. Rajmani, Nagasuma Chandra, Harinath Chakrapani, Karl Drlica, Amit Singh

**Author notes:** Equal Contribution (Somnath Shee, Samsher Singh). For correspondence (Amit Singh), (Karl Drlica).

## Abstract

Moxifloxacin is central to treatment of multidrug-resistant tuberculosis. Effects of moxifloxacin on *Mycobacterium tuberculosis* redox state were explored to identify strategies for increasing lethality and reducing the prevalence of extensively resistant tuberculosis. A non-invasive redox biosensor and an ROS-sensitive dye revealed that moxifloxacin induces oxidative stress correlated with *M. tuberculosis* death. Moxifloxacin lethality was mitigated by supplementing bacterial cultures with an ROS scavenger (thiourea), an iron chelator (bipyridyl), and, after drug removal, an antioxidant enzyme (catalase). Lethality was also reduced by hypoxia and nutrient starvation. Moxifloxacin increased the expression of genes involved in the oxidative stress response, iron-sulfur cluster biogenesis, and DNA repair. Surprisingly, and in contrast with *Escherichia coli* studies, moxifloxacin decreased expression of genes involved in respiration, suppressed oxygen consumption, increased the NADH/NAD^+^ ratio, and increased the labile iron pool in *M. tuberculosis*. Lowering the NADH/NAD^+^ ratio in *M. tuberculosis* revealed that NADH-reductive stress facilitates an iron-mediated ROS surge and moxifloxacin lethality. Treatment with N-acetyl cysteine (NAC) accelerated respiration and ROS production, increased moxifloxacin lethality, and lowered the mutant prevention concentration. Moxifloxacin induced redox stress in *M. tuberculosis* inside macrophages, and co-treatment with NAC potentiated the anti-mycobacterial efficacy of moxifloxacin during nutrient starvation, inside macrophages, and in mice where NAC restricted the emergence of resistance. Thus, oxidative stress, generated in a novel way, contributes to moxifloxacin-mediated killing of *M. tuberculosis.* The results open a way to make fluoroquinolones more effective anti-tuberculosis agents and provide a mechanistic basis for NAC-mediated enhancement of fluoroquinolone lethality *in vitro* and *in vivo*.

**Author Summary:** A new paradigm was revealed for stress-mediated bacterial death in which moxifloxacin treatment of *M. tuberculosis* decreases respiration rate (respiration <u>increases</u> in *E. coli*). Although moxifloxacin-induced, ROS-mediated bacterial death was observed, it derived from elevated levels of NADH and iron, a phenomenon not seen with antibiotic-treated *E*. *coli*. Nevertheless, stimulation of respiration and ROS by N-acetyl cysteine (NAC) enhanced moxifloxacin-mediated killing of *M. tuberculosis*, thereby reinforcing involvement of ROS in killing. NAC stimulation of moxifloxacin-mediated killing of *M. tuberculosis* and restriction of the emergence of resistance in a murine model of infection emphasize the importance of lethal action against pathogens. The work, plus published benefits of NAC to TB patients, encourage studies of NAC-based enhancement of fluoroquinolones.

## Introduction

Antimicrobial resistance is a growing problem for the management of tuberculosis (TB). For example, between 2009 and 2016, the number of global cases of multidrug-resistant tuberculosis (MDR-TB), defined as resistance of *Mycobacterium tuberculosis* to at least rifampicin and isoniazid, increased annually by over 20% (Lange et al., 2018). Additional resistance to a fluoroquinolone and at least one of three injectable agents (kanamycin, amikacin, or capreomycin), termed extensively drug-resistant tuberculosis (XDR-TB), accounted for about 6% of MDR-TB cases in 2018 (WHO, 2019). Alarmingly, the annual XDR-TB cases reported worldwide increased almost 10-fold between 2011 and 2018 (WHO, 2013; WHO, 2019). Increased prevalence of XDR-TB is not surprising, since the fluoroquinolone-containing combination therapies used to halt progression to XDR-TB utilized early quinolones (ciprofloxacin, ofloxacin, and sparfloxacin) that are only modestly effective anti-TB agents (Shandil et al., 2007).

Efforts to find more active fluoroquinolones led to C-8 methoxy derivatives that exhibit improved ability to kill *M. tuberculosis in vitro* (Dong et al., 1998). Two of these compounds, moxifloxacin and gatifloxacin, have been examined as anti-TB agents (Malik et al., 2006; Rodríguez et al., 2001; Ruan et al., 2016). Preclinical studies with moxifloxacin in murine TB models demonstrate effective treatment with reduced relapse frequency as well as treatment-shortening properties (Nuermberger et al., 2004). Indeed, single-drug clinical studies indicate that the early bactericidal activity of moxifloxacin is similar to that of first-line anti-TB-drugs, such as isoniazid and rifampicin. Thus, moxifloxacin could be part of effective multidrug combination treatments to shorten therapy time (Dorman et al., 2021; Pletz et al., 2004). Shorter treatment could increase treatment compliance, further limiting the progression of MDR- to XDR-TB. However, multiple phase III clinical trials have failed to corroborate the preclinical efficacy data, as moxifloxacin did not shorten treatment time (Gillespie et al., 2014; Jawahar et al., 2013; Jindani et al., 2014). One approach for increasing moxifloxacin efficacy is to find ways to increase its lethal action.

The fluoroquinolones have two mechanistically distinct anti-bacterial effects: 1) they block growth by forming reversible drug-gyrase-DNA complexes that rapidly inhibit DNA synthesis, and 2) they kill cells (Drlica and Zhao, 2020). Death appears to arise in two ways, one through chromosome fragmentation and the other through accumulation of reactive oxygen species (ROS) (Drlica and Zhao, 2020). The latter appear to dominate when DNA repair is proficient (Hong et al., 2020; Hong et al., 2019). We are particularly interested in fluoroquinolone lethality, because *M. tuberculosis* possesses a remarkable ability to evade host immune pressures (Ehrt and Schnappinger, 2009). In the absence of an effective immune response, bactericidal drugs are essential for clearing infection quickly. If the redox-based mechanisms of fluoroquinolone lethality seen with *E. coli* extend to *M. tuberculosis,* opportunities may exist for increasing ROS and lethality, thereby reducing treatment duration and increasing cure rate for MDR-TB.

Several methods are available for assessing the contribution of ROS to bactericidal activity. Among these are direct detection of ROS levels using ROS-sensitive dyes, characterization of mutants known to alter ROS levels, and examining effects of agents known to suppress ROS accumulation. The present work adds detection of antibiotic-induced changes in redox physiology of *M. tuberculosis* using a non-invasive, genetic biosensor (Mrx1-roGFP2) (Bhaskar et al., 2014). The sensor measures the redox potential (*E_MSH_*) of a major mycobacterial thiol buffer, mycothiol, thereby assessing oxidative effects independent of radical-sensitive fluorescent dye methods whose interpretation has been debated (Dwyer et al., 2014; Liu and Imlay, 2013). Mrx1-roGFP2 is well-suited for this work, as it has been used to measure *E_MSH_* of *M. tuberculosis* during *in vitro* growth, infection of macrophages, and exposure to non-quinolone antimicrobials (Bhaskar et al., 2014; Mishra et al., 2019; Mishra et al., 2017; Nambi et al., 2015).

In the present work, we began by asking how well *M. tuberculosis* fits the paradigm for ROS-mediated killing by antimicrobials as developed from studies using *E. coli*. We used moxifloxacin as a lethal stressor, because the fluoroquinolones are among the better understood agents and because moxifloxacin is potentially important for tuberculosis control. *M. tuberculosis* deviated markedly from *E. coli* by suppressing respiration rather than increasing it during fluoroquinolone-mediated stress. That raised questions about the source of ROS accumulation, since with *E. coli* ROS levels correlate with increased, not decreased, respiration. We found that an increase in NADH (reductive stress) accounted for the increase in ROS and lethality. When we artificially raised respiration, ROS levels increased, and we observed enhanced moxifloxacin lethality *in vitro,* inside macrophages, and in a murine model of infection. Raising respiration by N-acetyl cysteine (NAC) treatment also reduced the emergence of resistance. Thus, the work provides a mechanistic basis for developing respiration enhancers to increase the lethal action of moxifloxacin and perhaps other anti-TB agents.

## Results

### Moxifloxacin-mediated lethality associated with oxidative stress during aerobic growth

To obtain a reference point for *M*. *tuberculosis* susceptibility to moxifloxacin, we determined minimal inhibitory concentration (MIC) using the resazurin microtiter assay (REMA) (Padiadpu et al., 2016). MIC for moxifloxacin, which ranged from 0.125 μM to 0.5 μM, was similar for H37Rv and several drug-resistant isolates (Table S1; isolate similarity was also seen for levofloxacin and ciprofloxacin). Thus, strain H37Rv appeared to be representative and appropriate for subsequent experiments.

As an initial probe for moxifloxacin-induced oxidative stress, we examined a derivative of *M. tuberculosis* H37Rv that expresses the redox biosensor Mrx1-roGFP2 (strain *Mtb*-roGFP2). This biosensor reports the redox state of the mycothiol redox couple (reduced mycothiol [MSH]/oxidized mycothiol [MSSM]) in the bacterial cytosol (Fig 1A) (Bhaskar et al., 2014). Exposure of *Mtb*-roGFP2 to hydrogen peroxide (H_2_O_2_) led to rapid (5 min) oxidation of the biosensor in a concentration-dependent manner (Fig S1) in which a 2-fold increase in the biosensor ratio represented exposure to 500 μM of H_2_O_2_ (Fig S1).

**Fig 1.**
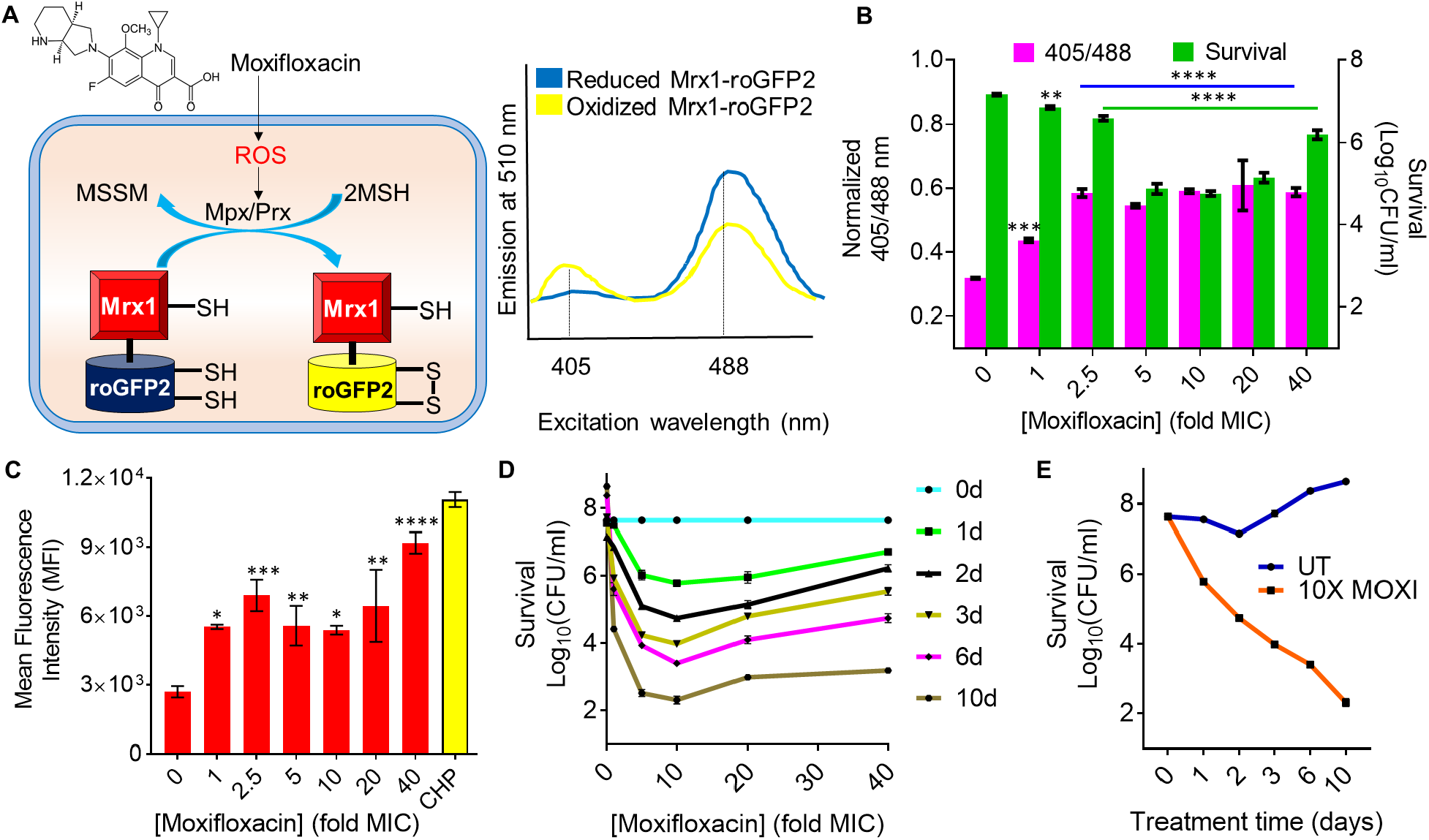
Effect of moxifloxacin on biosensor oxidation, ROS level, and bacterial survival. (**A**) Moxifloxacin increases oxidative stress that increases the ratio of oxidized (MSSM) to reduced mycothiol (MSH) via mycothiol-dependent peroxiredoxin (Prx) or peroxidase (Mpx). Oxidation of Mrx1-roGFP2 increases fluorescence intensity for excitation at ∼400 nm and decreases it for excitation at ∼490 nm. (**B**) Strain *M. tuberculosis*-roGFP2 was exposed to the indicated concentrations of moxifloxacin (1X MIC = 0.5 µM) for 48 h, and the ratiometric response of the biosensor and survival following treatment were determined. (**C**) Cells were treated as in panel B, and ROS were quantified by flow-cytometry using CellROX Deep Red dye. 10 mM cumene hydroperoxide (CHP) served as a positive control. Data represent mean fluorescence intensity of the dye. (**D**) Exponentially growing *M. tuberculosis* H37Rv was treated with moxifloxacin at the indicated concentrations for the indicated times; survival was assessed by determining colony-forming units (CFU). (**E**) Time-kill curves for *M. tuberculosis* treated with 10X MIC of moxifloxacin. Error bars represent standard deviation from the mean. Data represent at least two independent experiments performed in at least duplicate. Statistical significance is calculated against the no-treatment control (\*\*\*\**p* < 0.0001, *** *p* < 0.001, and \*\**p* < 0.01, \**p* < 0.05).

Increasing moxifloxacin concentration, up to 2.5X MIC (1X MIC = 0.5 µM) during a 48-h incubation, increased the biosensor ratiometric signal by almost 2-fold (Fig 1B, S2A, S2C), indicating elevated levels of oxidized mycothiol (MSSM). Above 2.5X MIC, MSSM levels appeared to plateau. Measurement of ROS with CellROX Deep Red showed a similar increase to 2.5X MIC moxifloxacin, followed by a plateau of mean fluorescence intensity (Fig 1C). Moxifloxacin treatment damaged DNA and oxidized lipids (Fig S3), indicating that drug-induced oxidative stress kills *M. tuberculosis* by degrading essential biomolecules.

Maximum bacterial killing was observed at 10X MIC, a concentration at which survival decreased by ≈ 2 log_10_-fold at treatment day 1 and by 6 log_10_-fold at treatment day 10 (Fig 1D, 1E). Survival dropped in a concentration-dependent manner, with a minimum being reached at a higher concentration than observed for maximal oxidative stress (10X MIC, Fig 1B, 1C, 1D). The discordance between the biosensor signal plateauing at 2.5X MIC and killing continuing to 10X MIC (Fig 1B) may derive from the existence of two lethal mechanisms. One mechanism, chromosome fragmentation (Dwyer et al., 2007; Malik et al., 2006), likely dominates at high quinolone concentration, with an ROS-based mechanism dominating at lower concentrations. Indeed, for norfloxacin treatment of *E. coli,* oxidative effects on killing are seen only at low-to-moderate drug concentrations (Malik et al., 2007). As expected, oxidative stress was not observed in *M. tuberculosis* treated with sub-inhibitory concentrations of moxifloxacin (Fig S2A). The increase in biosensor signal was observed as early as 12 h after initiating moxifloxacin treatment (half the doubling time of untreated control cells); it then increased significantly at 48 h (Fig S2A, S2B). Thus, ROS production precedes and contributes to the death of *M. tuberculosis.* As expected, ROS levels in a moxifloxacin-resistant isolate of *M. tuberculosis* remained low upon treatment with moxifloxacin (Fig S4A).

We also compared the ability of a weakly effective fluoroquinolone (ciprofloxacin, MBC = 2 μM; Table S2), a moderately effective compound (levofloxacin, MBC = 1 μM), and moxifloxacin (MBC = 0.5 μM) to oxidize the biosensor, all at 2.5-fold MBC. Moxifloxacin induced biosensor oxidation after treatment for 12 h and 24 h (Fig S4B). In contrast, ciprofloxacin did not induce biosensor oxidation, and levofloxacin triggered oxidative stress only after a 24-h treatment (Fig S4B). Thus, ROS levels do not correlate with MBC, a parameter obtained after long incubation time (see Discussion). The more active quinolones are also more lethal at the same fold MIC with several bacterial species, including mycobacteria (Dong et al., 1998; Zhao et al., 1998; Zhao et al., 1997), these data corroborate the link between oxidative stress and moxifloxacin lethality.

High-dose chemotherapy is postulated to slow the emergence of drug resistance if the drug concentration is above the mutant prevention concentration (MPC) (Drlica and Zhao, 2007). However bacterial strains can paradoxically show elevated survival levels at extremely high drug concentrations. In the case of nalidixic acid at very high concentration, *E. coli* survival can be 100% (Luan et al., 2018). Similarly, *M. tuberculosis* survival increased dramatically at high moxifloxacin concentration (Fig 1D).

Some aspects of high-concentration survival are discordant when *E. coli* and *M*. *tuberculosis* are compared. For example, with *E. coli* this phenotype is associated with reduced ROS levels (Luan et al., 2018); with *M*. *tuberculosis*, the biosensor and CellROX signals remained high at high levels of moxifloxacin (Fig 1B, 1C). This difference between the organisms, which is unexplained, encouraged further comparisions.

### Reduction of moxifloxacin-mediated killing of *M. tuberculosis* by ROS-mitigating agents

Work with *E. coli* supports the idea that ROS contribute to quinolone-mediated killing of bacteria (Drlica and Zhao, 2020). For example, superoxide (O_2_^.-^) damages Fe-S clusters and increases the free iron (Fe) pool (Keyer and Imlay, 1996), which may then drive the generation of toxic hydroxyl radical (HO.) via the Fenton reaction (Dwyer et al., 2007; Kohanski et al., 2007). To examine the effects of Fe, we assessed ROS levels in *M. tuberculosis* grown in minimal medium under Fe-deficient and Fe-excess conditions. Fe-overload raised ROS levels (Fig S5). Treatment with thiourea (TU), a thiol-based scavenger of ROS, then reversed the Fe-induced ROS increase (Fig S5). During moxifloxacin treatment of *M. tuberculosis*, a concentration-dependent increase in free Fe occurs (Fig S6). Pretreatment with thiourea, at the non-toxic concentration of 10 mM (Nandakumar et al., 2014), increased survival of *M. tuberculosis* by almost 10-fold during co-treatment with moxifloxacin (1X – 10X MIC) (Fig 2A, 2B, S7). These data support the connection between ROS accumulation and moxifloxacin-mediated killing.

**Fig 2.**
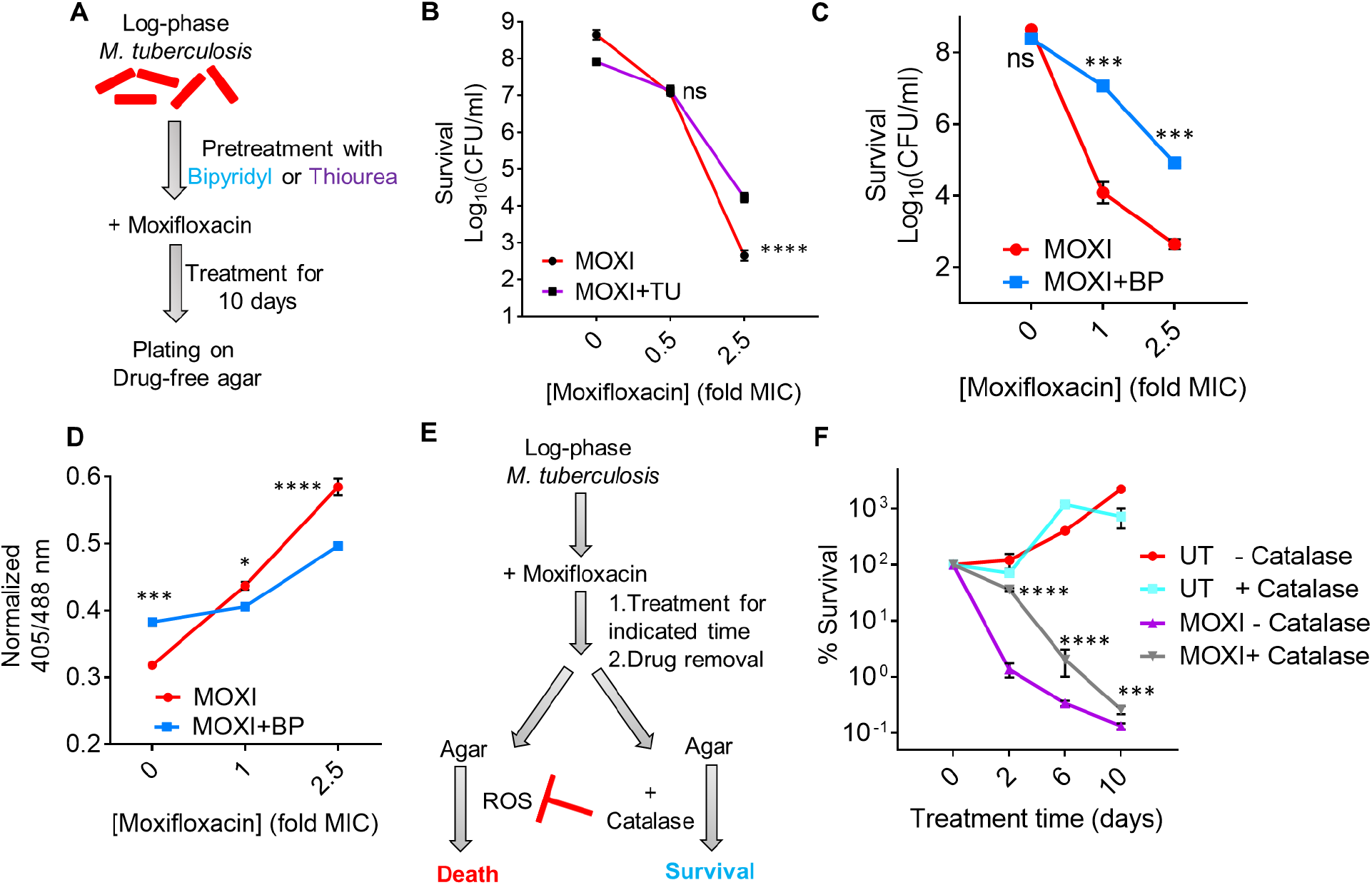
ROS mitigation reduces moxifloxacin-mediated killing of *M. tuberculosis*. (**A**) Plan for detecting thiourea (TU) and bipyridyl (BP) effects on moxifloxacin lethality. (**B**) Exponentially growing *M. tuberculosis* H37Rv cultures were either left untreated or treated with 10 mM TU for 1 h before addition of the indicated concentrations of moxifloxacin (MOXI; 1X MIC = 0.5 µM) for 10 days followed by determination of CFU. (**C**) Effect of bipyridyl. *M. tuberculosis* as in *B* was untreated or treated with 250 µM BP for 15 min prior to addition moxifloxacin as in *B*. (**D**) The Mrx1-roGFP2 biosensor ratiometric response was determined after 48 h treatment of *M. tuberculosis* cultures with the indicated concentrations of MOXI alone or with 250 µM of BP. (**E**) *M. tuberculosis* cultures were treated with moxifloxacin, the drug was removed by washing, and cells were plated on drug-free 7H11 agar with or without catalase followed by CFU determination. (**F**) *M. tuberculosis* cultures were treated with 1X MIC of moxifloxacin for the indicated times, washed, and plated with or without catalase (17.5 U/mL of agar). Percentage survival was calculated relative to CFU of cultures at 0 h. Statistical significance was calculated between drug-alone group with drug + catalase group. Statistical considerations were as in Fig 1.

Another test involves chelating free ferrous Fe with bipyridyl, a high-affinity Fe^2+^ chelator: with *E. coli*, bipyridyl lowers the lethal action of multiple lethal stressors (Hong et al., 2017; Kohanski et al., 2007). When we treated *M. tuberculosis* cultures with a non-inhibitory concentration (250 µM) of bipyridyl before moxifloxacin (Fig 2A), killing was reduced by >100 fold (Fig 2C), and ROS accumulation was lowered (Fig 2D). Treatment with bipyridyl or thiourea also protected from moxifloxacin-mediated DNA damage and lipid peroxidation (Fig S3A, S3B). Collectively, the data indicate that Fe-mediated ROS production is the underlying oxidative stress associated with moxifloxacin lethality.

Work with *E. coli* also indicates that ROS accumulates even after removal of a lethal stressor (Hong et al., 2019). When we treated *M. tuberculosis* with moxifloxacin in liquid medium and then plated cells on antibiotic-free 7H11 agar (see scheme, Fig 2E, we found that addition of catalase to the agar reduced bacterial killing (Fig 2F; peroxide diffuses freely across the cell membrane (Seaver and Imlay, 2001), allowing exogenous catalase to reduce endogenous ROS levels). Replacement of catalase with 2.5% bovine serum albumin, which is not expected to degrade peroxide, failed to protect *M. tuberculosis* from moxifloxacin-mediated killing (Fig S8). Finding that an anti-ROS agent suppresses killing after the removal of moxifloxacin indicates that the primary damage (rapid formation of fluoroquinolone-gyrase-DNA complexes) can be insufficient to kill *M. tuberculosis*. Thus, the post-stressor accumulation of ROS and death seen with *E. coli* occurs with *M. tuberculosis*.

We note that shorter drug-treatment times in 7H9 broth (2 days) prior to catalase treatment on drug-free agar demonstrated higher survival than longer times (6 to 10 days), but eventually there was little difference (Fig 2F). A similar phenomenon has been observed with *E. coli* (Hong et al., 2019). These data likely reflect ROS acting rapidly to cause accelerated lethality and then having little effect on long-term killing. Overall, studies with ROS-mitigating agents indicate that ROS contribute causally to quinolone-mediated lethality with both *E. coli* and *M. tuberculosis*.

### Reduction of moxifloxacin lethality by nutrient starvation and hypoxia

Our data support previous work (Gengenbacher et al., 2010) in which nutrient starvation blocked the lethal action of moxifloxacin (Loebel cidal concentration (LCC_90_) > 32 µM) (Fig S9A, S9B, Table S2). This phenomenon has also been reported for older quinolones with *E. coli* (Hong et al., 2020). With *E. coli*, these effects have been attributed to a metabolic shift that suppresses respiration, which is thought to be a major source of ROS.

Previous work (Gengenbacher et al., 2010; Wayne and Hayes, 1996) also shows that hypoxic conditions reduce the lethal action of older fluoroquinolones with *M. tuberculosis*. We confirmed this observation by showing that moxifloxacin-mediated killing decreases significantly under hypoxic conditions (Fig S9A, S9C, Table S2). With the Wayne Cidal Concentration (WCC_90_) assay (Gengenbacher et al., 2010), the moxifloxacin concentration required to kill 90% of hypoxic bacteria after 5 days of drug treatment was ∼10 µM (Table S2). This value corresponds to 20X MIC or 22-fold higher than the LD_90_ (lethal dose) with *M. tuberculosis* (0.45 µM) cultured under aerobic growth conditions for the same treatment time (Fig S10).

The hypoxia findings fit with moxifloxacin lethality being associated with an increase in ROS, an event that should be suppessed by O_2_ limitation (intracellular generation of ROS may depend largely on the transfer of electrons directly to molecular O_2_ (Imlay, 2013)). However, hypoxic suppression of moxifloxacin lethality could be due to decreased respiration. To test this idea, we stimulated anaerobic respiration by providing nitrate as an alternative electron acceptor (Wayne and Hayes, 1998), since that restores norfloxacin lethality with anaerobic *E. coli* (Dwyer et al., 2014). Surprisingly, nitrate lowered residual anaerobic lethality of moxifloxacin by several fold (Fig S9D). Nitrate was even more protective with *M. tuberculosis* exposed to metronidazole (Fig S9D), an antimicrobial known to be lethal in the absence of oxygen (Wayne and Sramek, 1994). The surprising protective effect of nitrate, which is considered in more detail in the Discussion, encouraged further comparisons with *E. coli*.

### Effect of moxifloxacin on the *M. tuberculosis* transcriptome

When we examined the *M. tuberculosis* transcriptome using published data from a 16-h exposure to 2X, 4X, and 8X MIC moxifloxacin (Ma et al., 2015), we found that 359 genes exhibited altered expression (2-fold change across all three treatment conditions). Of these, 219 genes were upregulated, and 140 were down-regulated (Dataset S1). We asked whether expression after prolonged moxifloxacin exposure (16 h) is largely a secondary effect from induction of other genes by comparing early changes (4-h moxifloxacin exposure) in expression for a set of 28 genes deregulated at 16 h. We saw little difference (Fig S11A, S11B). Thus, the patterns we observed at 16 h appear to largely reflect a primary transcription response.

When we classified the differentially expressed genes (DEGs) according to annotated functional categories (Kapopoulou et al., 2011), we found that “Information Pathways” and “Insertion Sequences and Phages” were two-fold over-represented in the moxifloxacin-treated *M. tuberculosis* transcriptome (Table S3). These data suggest that the bacterium responds to the drug mainly by regulating DNA remodeling, transcription, and translational machinery.

Increased expression was also seen with genes involved in redox homeostasis, such as thioredoxin (*trxB1, trxB2,* and *trxC*), alkyl hydroperoxide reductase (*ahpC*), the SigH/RshA system, the copper-sensing transcriptional regulator *csoR,* and the *whiB*-family (*whiB4* and *whiB7*) (Fig 3A). Indeed, the transcriptome of moxifloxacin-treated *M. tuberculosis* showed 67% overlap with the transcriptional signature of H_2_O_2_-treated cells (Fig 3B) (Voskuil et al., 2011). Statistical analysis showed a signficant overlap between the moxifloxacin transcriptome and the response to oxidative stress (H_2_O_2_; *p* = 5.39 e-14) and nitrosative stress (NO; *p* = 2.13 e-8) (Fig S12, Table S4). An induction of oxidant-responsive genes (*sox, sod, mar*) is also seen in norfloxacin-treated *E. coli* (Dwyer et al., 2007). Other conditions, such as hypoxia and acidic pH, showed non-significant overlap with the moxifloxacin transcriptome (Fig S12, Table S4). Collectively, the data are consistent with the idea that fluoroquinolones stimulate ROS accumulation.

**Fig 3.**
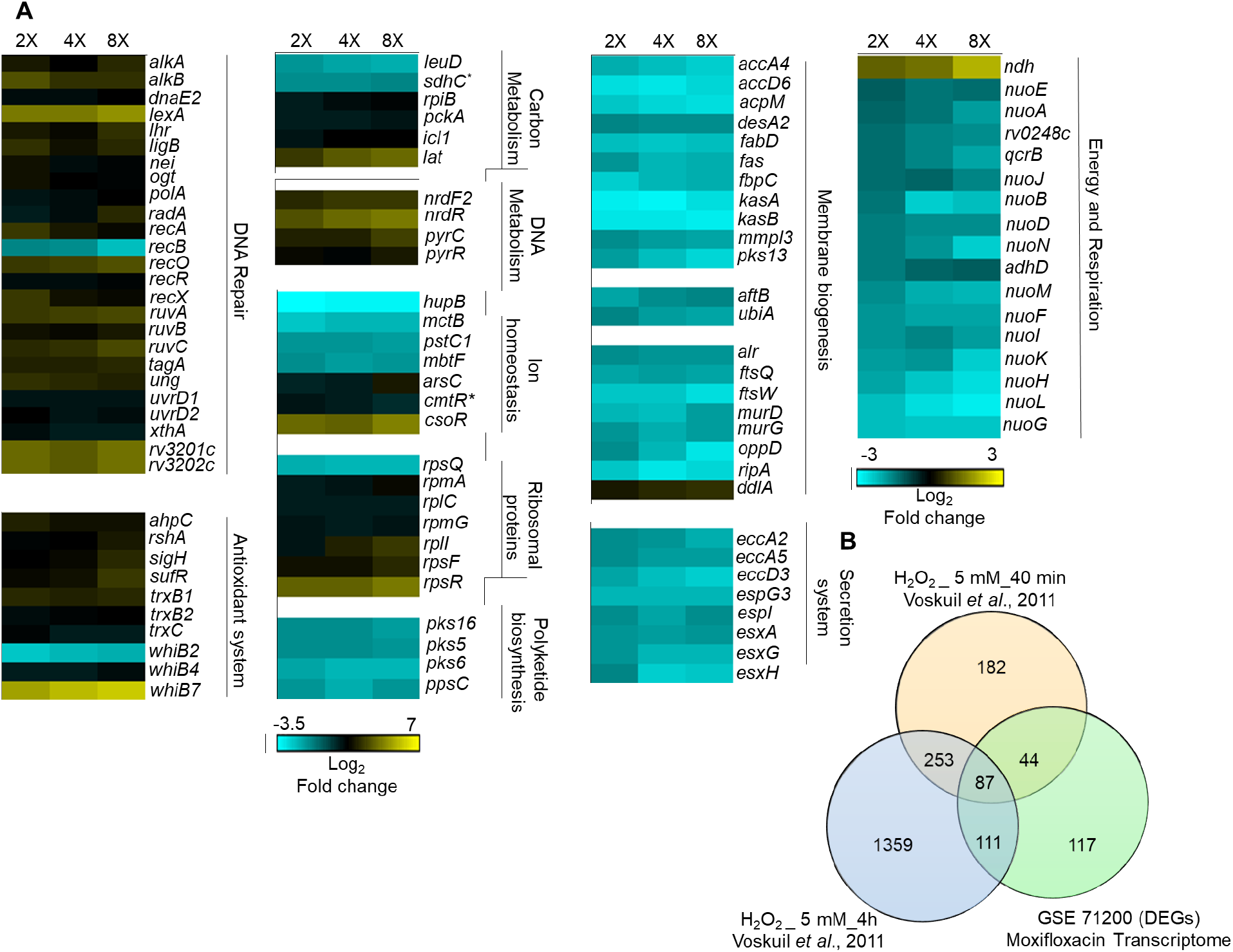
Whole-genome transcriptome profiling of *M. tuberculosis* treated with moxifloxacin. (**A**) Heat-map showing expression changes due to 16-h treatment of *M. tuberculosis* with moxifloxacin. Differentially expressed genes (DEGs) exhibited a 2-fold change across all three treatment conditions (2X, 4X, and 8X MIC of moxifloxacin; 1X MIC = 0.4 µM); color code for the fold change is at the bottom of the second column (yellow: upregulated genes; turquois: down-regulated genes). Genes are grouped according to function. For genes belonging to bioenergetics processes, color code for the fold change is at the bottom of the fourth column. \**sdhC* and *cmtR* are deregulated in two treatment conditions (2X and 4X) (**B**) Venn diagram showing transcriptome overlap between moxifloxacin-mediated (green circle) and H_2_O_2_-mediated stress for *M. tuberculosis* (Voskuil et al., 2011); DEGs obtained with treatment with 5 mM H_2_O_2_ for 40 min and 5 mM H_2_O_2_ for 4 h are shown in beige and blue circles, respectively.

Several genes involved in repairing DNA (*recA/O/R/X, ruv ABC, uvrD, ung, dnaE2, xthA, radA, alkA, lexA, nei, ligB*), in DNA metabolism (*nrdF2, nrdR, pyrC, pyrR*), and in Fe-S cluster biogenesis (*sufR*) (Anand et al., 2021; Das et al., 2021) were also upregulated (Fig 3A). *E. coli* treated with norfloxacin shows similar upregulation of DNA-damage responses (*lexA, uvr, rec* systems and error-prone DNA polymerases IV and V), nucleotide metabolism, and Fe-S cluster biogenesis (*IscRUSA*) (Dwyer et al., 2007). Such results are expected from the known ability of ROS to damage DNA (Dwyer et al., 2015; Foti et al., 2012).

With *M. tuberculosis*, moxifloxacin suppressed expression of an Fe-responsive repressor (*hupB*) and an iron siderophore (*mbtF*), consistent with drug treatment increasing the intrabacterial pool of labile Fe (Fig S6). Additionally, the microarray data suggest that Fe–S clusters in *M. tuberculosis* are exposed to ROS during treatment with moxifloxacin, resulting in Fe release from damaged clusters and increased expression of Fe-S repair pathways (*sufR*). In contrast, *E. coli* upregulates Fe-uptake machinery (Dwyer et al., 2007). Thus, *M. tuberculosis* and *E. coli* may generate ROS in different ways.

In *M. tuberculosis*, moxifloxacin repressed the expression of energy-efficient respiratory complexes, such as succinate dehydrogenase (*rv0248c, sdhC*), cytochrome bc1 (*qcrB*), and type I NADH dehydrogenase (*nuo* operon) (Fig 3A). In contrast, the level of the energetically inefficient, non-proton pumping *ndh2* was increased. These data indicate that *M. tuberculosis* slows primary respiration and shifts to a lower energy state in response to moxifloxacin. Indeed, several energy-requiring pathways (*e.g.,* cell wall biosynthesis, cofactor biogenesis, cell division, transport, and ESX-secretion systems) were downregulated (Fig 3A and Dataset S1). Moreover, genes coordinating alternate carbon metabolism, such as the gluconeogenesis (*pckA*) and glyoxylate cycle (*icl1*), were up-regulated (Fig 3A). Interestingly, Icl1 protects *M. tuberculosis* from anti-TB drugs (isoniazid, rifampicin, and streptomycin), presumably by counteracting redox imbalance induced by these antibiotics (Nandakumar et al., 2014). The key idea is that *M. tuberculosis* enters into a quasi-quiescent metabolic state in response to stress from moxifloxacin, opposite to fluoroquinolone effects with *E. coli* (Dwyer et al., 2014; Dwyer et al., 2007).

### Moxifloxacin slows respiration

As a direct test for decelerated respiration, we measured the Extracellular Acidification Rate (ECAR) and Oxygen Consumption Rate (OCR) for moxifloxacin-treated *M. tuberculosis* as readouts for proton-extrusion into the extracellular medium (reflecting glycolysis and TCA cycle activity) and for oxidative phosphorylation (OXPHOS), respectively, using a Seahorse XFp Analyzer (Lamprecht et al., 2016; Mishra et al., 2019). We incubated *M. tuberculosis* in unbuffered 7H9 + glucose medium in an XF microchamber, exposed the culture to 10X MIC moxifloxacin, and later to the uncoupler CCCP (addition of CCCP stimulates respiration to the maximal level). The difference between basal and CCCP-induced OCR estimates the spare (reserve) respiratory capacity (SRC) available to counteract stressful conditions (Lamprecht et al., 2016; Saini et al., 2016).

As expected for a growing, drug-free culture of *M. tuberculosis,* OCR showed a gradual increase over the duration of the experiment (400 min); OCR increased further upon uncoupling by CCCP (Fig 4A). In contrast, addition of 10X MIC of moxifloxacin inhibited the time-dependent increase in basal OCR (Fig 4A), and the level of OCR upon uncoupling by CCCP was markedly lower than in the absence of moxifloxacin (Fig 4A). In parallel, we measured viability of *M. tuberculosis* and found that the bacterial culture maintained 100% survival during the the entire incubation period and for an additional 4 h (Fig S13). Thus, the stalled oxygen-consumption effect of moxifloxacin cannot be attributed to dead cells.

**Fig 4.**
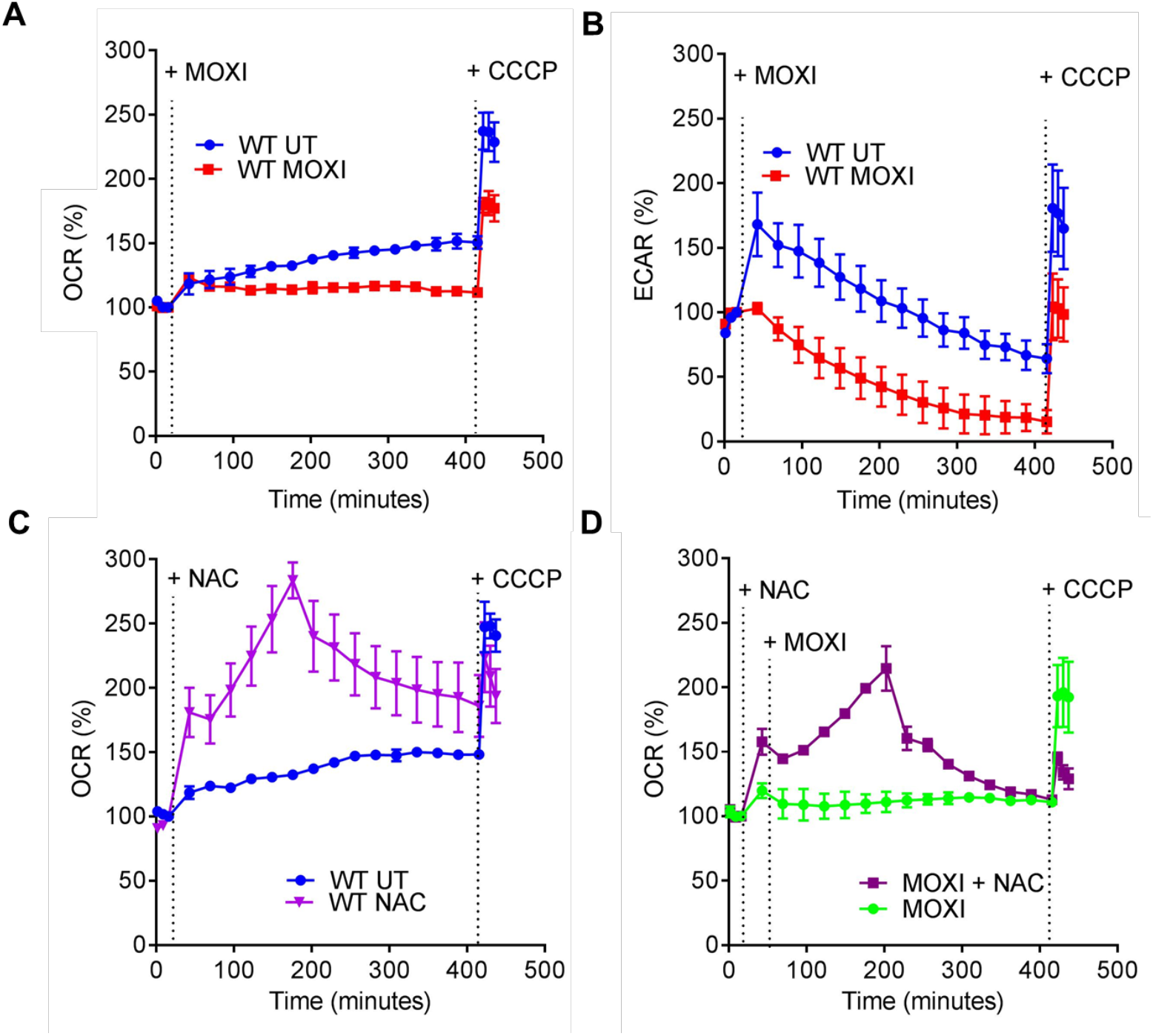
Moxifloxacin-mediated respiration arrest reversed by NAC. (**A**) OCR (pmol/min) indicated oxygen consumption rate. Exponentially growing *M. tuberculosis* cultures were either left untreated (UT) or treated with 10X MIC of moxifloxacin (MOXI; 5 µM) for the indicated times; black dotted lines indicate when MOXI or NAC (1 mM) or CCCP (10 µM) were added. Determination was via Seahorse XFp Analyzer (**B**) ECAR (mpH/min) indicated H^+^ production or extracellular acidification due to glycolytic and TCA flux. Determination was as in OCR with data representing percentage of third baseline value. NAC (1 mM) addition enhanced OCR of (**C**) untreated and (**D**) MOXI-treated cells. Data shown are representative of two independent experiments performed in triplicate.

The ECAR response increased for 60 min and then gradually dropped for untreated bacteria. This transient, early increase in ECAR was absent in cultures of moxifloxacin-treated *M. tuberculosis*, which exhibited a greater reduction of ECAR than untreated cultures at late times (Fig 4B). At the end of the experiment, moxifloxacin-treated cells were completely exhausted of glycolytic capacity (Fig 4B). The slowing in OCR and ECAR was also seen with treatment at lower concentrations of moxifloxacin (1X and 2.5X MIC) (Fig S14), while an anti-TB drug (ethambutol) that does not generate ROS, failed to significantly affect either OCR or ECAR (Fig S15).

In summary, moxifloxacin decelerates respiration and carbon catabolism in *M. tuberculosis*, which likely renders the bacterium metabolically quiescent and therefore less readily killed by moxifloxacin. In support of this idea, a recent study shows that pretreatment with the anti-TB drugs bedaquiline (an ATP synthase inhibitor) or Q203 (a cytochrome C oxidase inhibitor) reduces moxifloxacin-mediated killing of *M. tuberculosis* (Lee et al., 2019).

### ROS derive from increased NADH

Since moxifloxacin slows respiratory metabolism (Fig 4A) while increasing ROS levels (Fig 1B, 1C), a source of ROS other than respiration must exist. Slowed respiration during anoxia or chemical hypoxia places the electron transport chain (ETC) in a reduced state, and NADH accumulates. This phenomenon is known as reductive stress (Mavi et al., 2019). Using a redox cycling assay, we detected an accumulation of NADH in 1X, 2.5X and 10X MIC moxifloxacin-treated *M. tuberculosis* at 48 h (Fig 5A). NAD^+^ was raised at 1X MIC moxifloxacin, but it showed no significant difference from the untreated control at lethal, elevated moxifloxacin concentrations (Fig 5B); the total NAD/H pool decreased at high drug concentrations (2.5X and 10X MIC; Fig S16). Thus, the NADH/NAD^+^ ratio increased as moxifloacin concentration increased (Fig 5C), indicating reductive stress.

**Fig 5.**
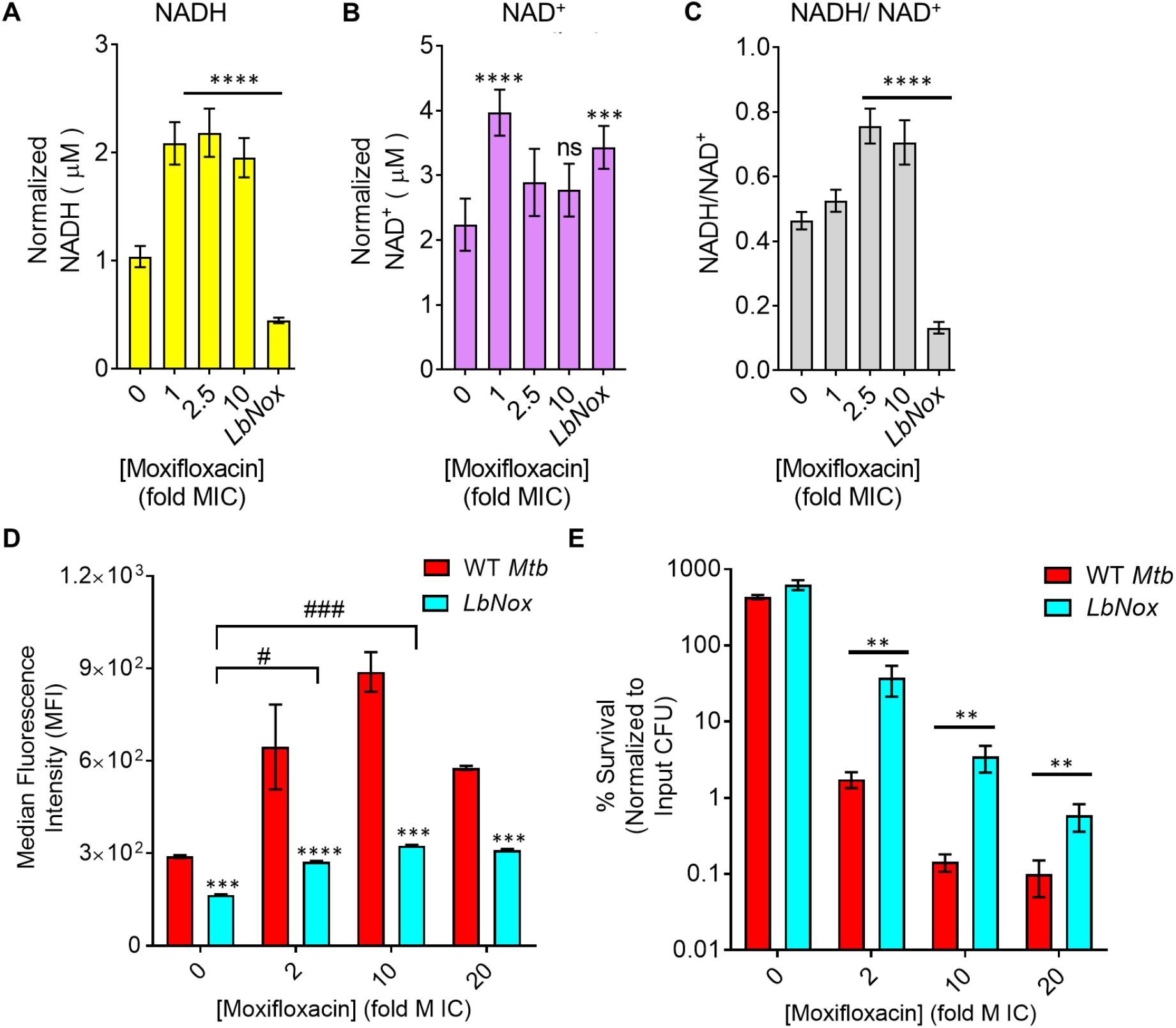
Dissipation of NADH reductive stress diminishes moxifloxacin-induced ROS increase and lethality with *M. tuberculosis.* Detection of **(A)** NADH or **(B)** NAD^+^ levels. *M. tuberculosis* was treated with moxifloxacin (1X MIC = 0.5 µM), for 2 days, and NADH or NAD^+^ levels were determined by an alcohol dehydrogenase-based redox cycling assay. **(C)** NADH/NAD^+^ ratio. Untreated *M. tuberculosis* expressing *LbNox* served as a control. *p* was determined by unpaired two-tailed student’s t-test analyzed relative to the untreated control. **(D)** ROS response to moxifloxacin. Wild-type *M. tuberculosis* H37Rv (WT *Mtb*) and cells expressing *Lbnox* (*LbNox*) were exposed to the indicated concentrations of moxifloxacin for 48 h, and ROS were quantified by flow-cytometry using CellROX Deep Red. **(E)** Cultures of exponentially growing wild-type *M. tuberculosis (*WT *Mtb)* and *LbNox* were treated with the indicated concentrations of moxifloxacin for 48 h, and survival was assessed by determining CFU. Statistical coniderations were as in Fig 1 (**** *p*< 0.0001, *** and ### *p* < 0.001, ** *p* < 0.01, # *p* < 0.05, ns indicates not significant).

We also measured NADH-reductive stress by expressing a genetically encoded, non-invasive biosensor of the NADH/NAD^+^ ratio (Peredox) (Bhat et al., 2016) (Fig S17). A ratiometric increase in Peredox fluorescence indicated an increase in the NADH/NAD^+^ ratio upon moxifloxacin treatment (1X to 40X MIC) at 48 h. We observed a similar increase in the Peredox ratio upon treatment of *M. tuberculosis* with bedaquiline, a drug known to increase the NADH/NAD^+^ ratio (Fig S17) (Bhat et al., 2016). As with NADH, NADPH accumulated, and the NADPH/NADP^+^ ratio increased in response to moxifloxacin treatment (Fig S18). Thus, moxifloxacin preferentially triggers NAD(P)H accumulation in *M. tuberculosis*.

In principle, stalled ETC and excessive reducing equivalents (NADH) can stimulate ROS production in two ways. In one, NADH mobilizes bound iron, thereby increasing the labile iron pool (Jaeschke et al., 1992). As previously mentioned, we found a drug-concentration-dependent increase in the free iron pool in *M. tuberculosis* after moxifloxacin treatment (Fig S6). Additionally, NADH autooxidation keeps iron in a reduced state (Fe^2+^); thus, NADH can drive the Fenton reaction towards hydroxyl radical generation (Jaeschke et al., 1992). Second, reduced components of the electron transport chain can directly transfer electrons to molecular oxygen to generate ROS (Kareyeva et al., 2012; Vinogradov and Grivennikova, 2016). If these ideas are correct, dissipation of the NADH overload, i.e., reductive stress, should lower the labile iron pool, ROS surge, and moxifloxacin lethality.

We lowered the NADH/NAD^+^ ratio with an *M. tuberculosis* strain that constitutively expresses *Lactobacillus brevis* NADH oxidase (*Mtb-LbNox*) (Fig 5C, S17) (Titov et al., 2016). In comparison to wild-type *M. tuberculosis*, *Mtb-LbNox* shows decreased levels of NADH and increased NAD^+^ without affecting the total pool of NAD^+^ + NADH (Fig 5A, 5B, 5C, S19). As expected, moxifloxacin-treated *Mtb-LbNox* displayed a significantly reduced free iron pool (Fig S6), ROS surge (Fig 5D), decreased DNA damage (Fig S6), and a 25-to 30-fold higher surivival than wild-type cells (Fig 5E). Thus, increased NADH/NAD^+^ ratio occurring during metabolic quiescence caused by moxifloxacin accounts for the increased labile iron pool, which in turn catalyzes Fenton-mediated production of ROS and cell death.

The growth, OCR, and ECAR of *Mtb-LbNox* in 7H9 broth were the same as with wild-type cells (Fig S20). Furthermore, MIC of moxifloxacin and other anti-TB drugs (rifampicin, ethambutol, and bedaquiline) with *Mtb-LbNox* was similar to that of wild-type *M. tuberculosis* (Table S1, S5), indicating that the interaction between drug and its primary target is unaffected by overexpression of *LbNox.* In contrast, *Mtb-LbNox* exhibited a 2-fold lower MIC for isoniazid, which is consistent with the interaction of isoniazid and its primary target (enoyl-ACP reductase, InhA) depending on the NADH/NAD^+^ ratio in *M. tuberculosis* (Vilchèze et al., 2005). Thus, overexpression of *LbNox* appears to have no effect on growth, metabolism, or respiration of *M. tuberculosis* that would complicate our interpretation that NADH dissipation protects from moxifloxacin-mediated killing.

### N-acetyl cysteine stimulates respiration, oxidative stress, and moxifloxacin lethality

Previous work showed that stimulating respiration using cysteine and N-acetyl cysteine (NAC) elevates the killing activity of combinations of anti-TB drugs (Vilchèze et al., 2017; Vilchèze and Jacobs, 2021). Since *M. tuberculosis* responded to moxifloxacin by dampening OXPHOS, we expected that countering the dampening with NAC would provide a way to increase ROS further and therefore enhance moxifloxacin lethality. We first measured time-dependent changes in OCR of *M. tuberculosis* upon exposure to a non-toxic dose of NAC (1 mM). Addition of NAC alone produced a sharp increase in OCR that reached its maximal level by 200 min (Fig 4C). At later times, OCR gradually declined; it did not increase upon addition of CCCP (Fig 4C), indicating that NAC-stimulated respiration exhausted the spare respiratory capacity of *M. tuberculosis*. Pre-treatment with NAC reversed moxifloxacin-mediated slowing of respiration (Fig 4D). As with NAC alone, the OCR of *M. tuberculosis,* treated with NAC plus moxifloxacin, increased for 200 min and then gradually dropped; CCCP remained ineffective at stimulating OCR (Fig 4D). These data indicate that NAC-stimulated oxygen consumption by *M. tuberculosis* overcomes the dampening effect of moxifloxacin on bacterial respiration.

Respiration, stimulated by NAC, augmented ROS accumulation upon moxifloxacin treatment and increased lethality (Fig 6A). NAC elevated the Mrx1-roGFP2-dependent redox signal for moxifloxacin-treated *M. tuberculosis*, confirming the increase in oxidative stress (Fig 6B, S2C, S2D). We emphasize that NAC had <u>no</u> effect on moxifloxacin MIC, as measured by REMA and 7H11 agar plate assay (Fig S21A, S21B); thus, the primary interaction between drug and DNA gyrase (cleaved complex formation) is unaffected by NAC. Supplementation of moxifloxacin at 1X and 5X MIC with 1 mM of NAC reduced survival by 21- and 11-fold, respectively (Fig 6C), levels that are consistent with recently published results using drug combinations (Vilchèze and Jacobs, 2021).

**Fig 6.**
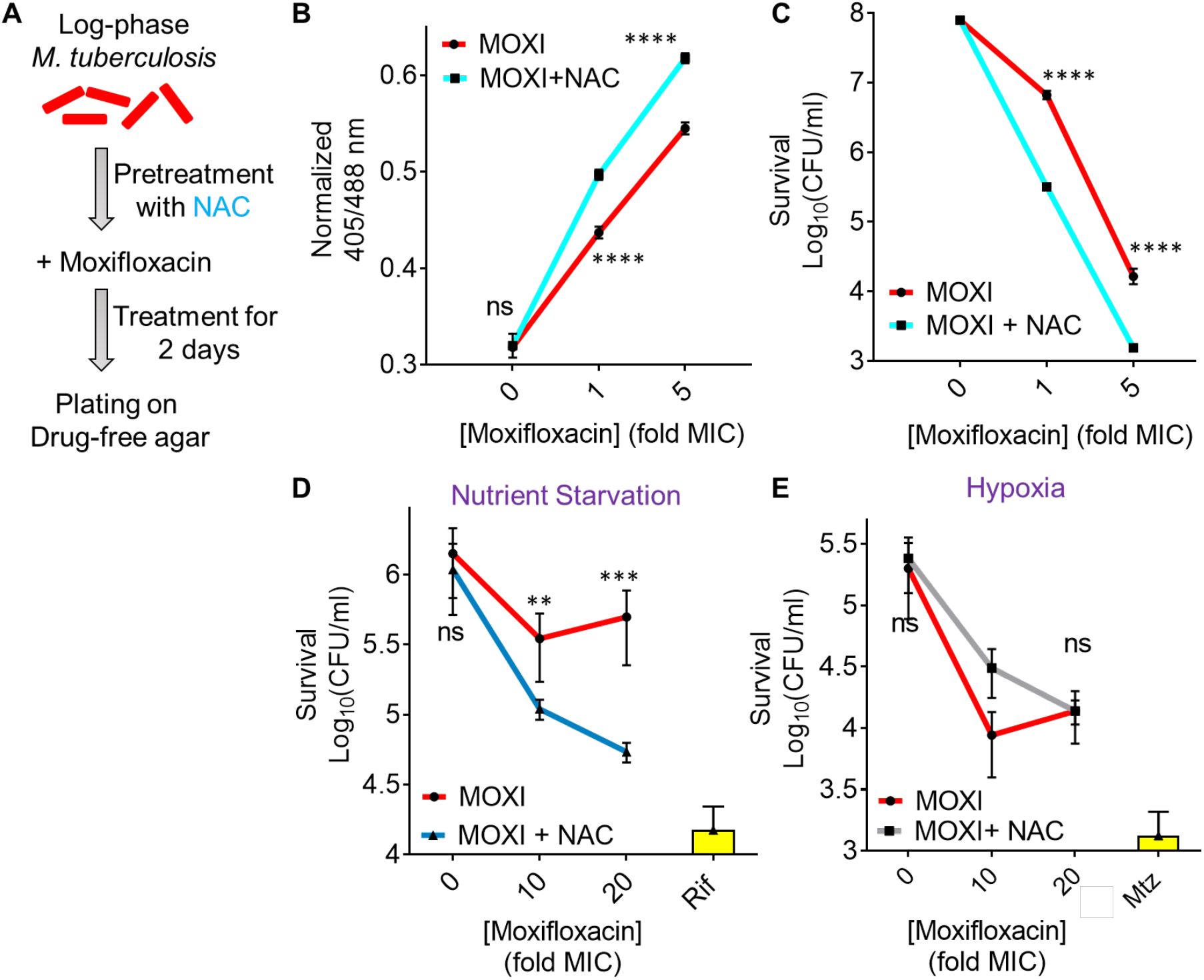
N-acetyl cysteine increases oxidative stress and moxifloxacin-mediated killing of *M. tuberculosis*. NAC (1 mM) was administered 1 h before addition of moxifloxacin (MOXI; 1X MIC = 0.5 µM) at the indicated concentrations followed by 48 h incubation. **(A)** Experimental plan. (**B**) Oxidative stress, measured by the ratiometric response of the Mrx1-roGFP2 biosensor. (**C**) Bacterial survival, measured by plating on 7H11 agar. **(D)** Effect of NAC and moxifloxacin combination with dormant bacilli. *M. tuberculosis* cultures were starved for nutrients for 14 days and then treated with moxifloxacin for 5 days in the presence or absence of NAC (1 mM) before determination of survival. Rifampicin (25 µM) served as a positive control. **(E)** NAC (1 mM) was added when cultures were placed in Vacutainer tubes followed by the treatment conditions indicated in Fig S2A and S2D. Metronidazole (Mtz; 10 mM) served as a positive control. Statistical considerations were as in Fig 1.

As a complement to the suppression of killing by addition of catalase after drug removal (Fig 2F), we treated *M. tuberculosis* cultures with 1X and 5X MIC of moxifloxacin and then plated the cells on 7H11 agar with 1 mM NAC. Post-drug addition of NAC blocked formation of colonies that would otherwise have been observed (Fig S22). Extended incubation (>4 weeks) resulted in the appearance of small colonies on the NAC-containing plates, which was likely due to instability/oxidation of NAC (T_1/2_∼ 14 h) under aerobic conditions (Held and Biaglow, 1994; Jaworska et al., 1999; Sommer et al., 2020).

NAC also potentiated moxifloxacin lethality (5- to 7-fold) with nutrient-starved, dormant *M. tuberculosis* (Fig 6D). These data agree with recent results in which activation of oxidative metabolism and respiration, through addition of L-cysteine to nutrient-starved cultures, reduced the fraction of persister subpopulations and enhanced the *in vitro* lethal activity of isoniazid and rifampicin (Srinivas et al., 2020). As expected, a similar treatment with NAC failed to enhance moxifloxacin lethality under hypoxic conditions (Fig 6E), consistent with NAC functioning by inducing oxygen consumption and elevating ROS levels.

To address the effect of NAC on the acquisition of moxifloxacin resistance, we measured the mutant prevention concentration (MPC) i.e., the minimal concentration of moxifloxacin at which no moxifloxacin-resistant clone emerges on moxifloxacin-containing 7H11 agar inoculated with ∼ 2.5 x 10^9^ bacteria (Drlica and Zhao, 2007; Singh et al., 2017). With a clinical MDR isolate of *M. tuberculosis* (NHN1664), moxifloxacin exhibited an MPC of 4 μM that was reduced to 2 µM upon co-treatment with either 1 mM or 2 mM NAC. For a lower moxifloxacin concentration (1 µM, 2X MIC), the fraction of cells recovered decreased by 10-fold and 100-fold when co-plated with 1 mM and 2 mM NAC, respectively (Table S6). Reduction of MPC is expected when lethal activity is increased (Cui et al., 2006).

We also addressed the possibility that some effects of NAC derived from adducts formed with moxifloxacin. A series of biochemical tests (thin layer chromatography, NMR, fluorescence assays, and LC-MS) revealed no evidence for adduct formation (Fig S23).

### Moxifloxacin-induced redox imbalance during infection of macrophages

The Mrx1-roGFP2 biosensor previously showed that first-line anti-TB drugs cause an oxidative shift in the *E_MSH_* of *M. tuberculosis* residing inside macrophages (Bhaskar et al., 2014). When we infected macrophages with *M. tuberculosis* H37Rv or an MDR clinical isolate (strain NHN1664), each expressing the biosensor Mrx1-roGFP2, *M. tuberculosis* displayed redox-heterogeneity (*E_MSH_*-basal [-280 mV], *E_MSH_-*oxidized [-240 mV], and *E_MSH_*-reduced [-320 mV]); the *E_MSH_*-reduced subpopulation was predominant (Bhaskar et al., 2014; Mishra et al., 2019; Mishra et al., 2021) (Fig 7A, S24). Treatment of *M. tuberculosis-*infected THP-1 macrophages with various concentrations of moxifloxacin significantly increased the *E_MSH_*-oxidized subpopulation (fuchsia colored line, Fig 7A, S24).

**Fig 7.**
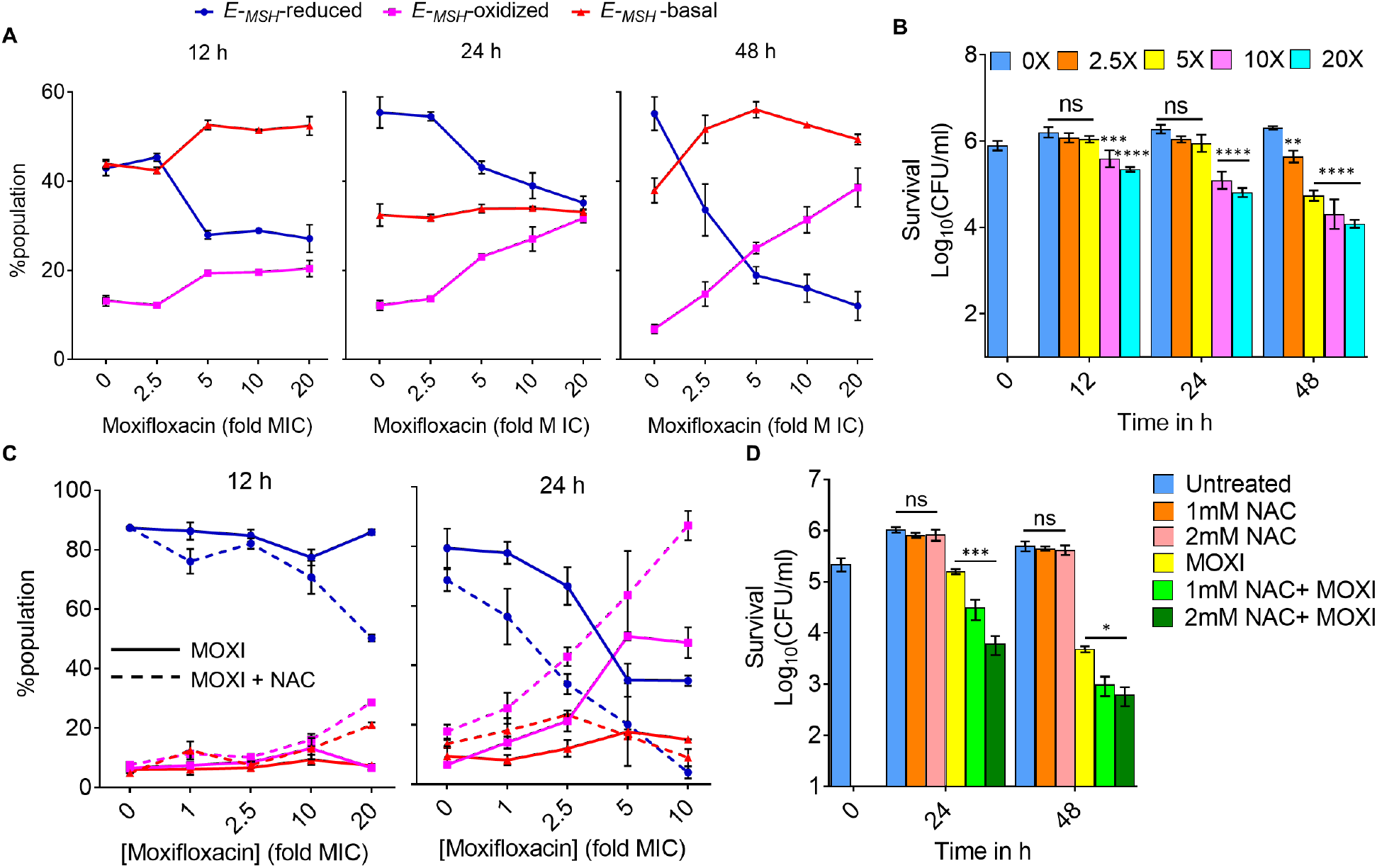
Moxifloxacin-induced oxidative shift in *E_MSH_* and killing of *M. tuberculosis* inside macrophages. THP-1 macrophages, infected with *Mtb*-roGFP2 (MOI = 1:10), were treated with moxifloxacin (MOXI; 1X MIC = 0.5 µM) immediately after infection and incubated for the indicated times. (**A**) ∼10,000 infected macrophages were analyzed by flow cytometry to quantify changes in the *E_MSH_* of *M. tuberculosis* subpopulations. (**B**) Bacterial survival kinetics after MOXI treatment of THP-1 macrophages infected with *Mtb*-roGFP2 (CFU determination). (**C**) *Mtb*-roGFP2-infected THP-1 macrophages were treated with MOXI at the indicated concentrations in presence or absence of NAC (1 mM) immediately after infection and incubated for the indicated times; analysis was as in *A.* (**D**) THP1 macrophages, infected by *Mtb*-roGFP2, were treated with NAC (1 mM or 2 mM), MOXI (10 μM), or the combination of NAC plus MOXI at those concentrations. After the indicated incubation times, the bacterial load in the macrophages was determined by plating on drug-free agar. *p* was determined by two-tailed student’s t-test compared to MOXI-alone treatment at each time-point. Statistical considerations were as in Fig 1.

The shift toward the *E_MSH_*-oxidized redox state was seen with moderate concentrations of moxifloxacin (5X MIC) at an early time point (12 h post infection), before *M. tuberculosis* survival started to drop (Fig 7A, 7B). Thus, moxifloxacin-mediated oxidative stress precedes killing: oxidative stress is not simply a consequence of cell death. Bacterial survival was reduced more by higher moxifloxacin concentrations (10X and 20X MIC) or longer incubation times (24 h and 48 h) (Fig 7B).

We also assessed the effect of NAC on *E_MSH_* and killing of *M. tuberculosis* inside macrophages. NAC itself is not cytotoxic to macrophages up to at least 5-10 mM (Amaral et al., 2016; Vilchèze et al., 2017), but at those elevated concentrations it exhibits anti-mycobacterial properties (Amaral et al., 2016; Cao et al., 2018). Consequently, we selected non-cytotoxic, non-anti-tuberculous concentrations of NAC (1 mM and 2 mM) (Amaral et al., 2016) for our test with moxifloxacin. NAC supplementation induced an oxidative shift in the *E_MSH_* of *M. tuberculosis* that exceeded that seen with moxifloxacin alone (Fig 7C, S24). NAC alone had no effect on survival of *M. tuberculosis* inside THP-1 macrophages; however, the NAC plus moxifloxacin combination decreased bacterial burden 5-10 times more than moxifloxacin alone in both concentration-and time-dependent manners (Fig 7D). We note that for either isoniazid or an isoniazid-rifampicin combination, NAC potentiates lethality only at late times (day 6 or 7) (Vilchèze et al., 2017; Vilchèze and Jacobs, 2021); NAC acts more rapidly with moxifloxacin, as the killing effect is evident at days 1 and 2 (Fig 7D). Thus, NAC may not function in the same way with all anti-TB drugs inside macrophages.

Moxifloxacin also lowered the level of an *E_MSH_*-reduced subpopulation inside macrophages (blue line, Fig 7A, 7C, S24); this subpopulation was further diminished by NAC supplementation. *E_MSH_*-reduced subpopulations are phenotypically tolerant to anti-TB drugs (Bhaskar et al., 2014; Mishra et al., 2019) and to moxifloxacin (Fig S25). Overall, results with infected macrophages demonstrate that accelerating respiration and oxidative metabolism by NAC can enhance moxifloxacin lethality.

### Potentiation of moxifloxacin lethality with NAC lowers *M. tuberculosis* burden in a murine model of infection and restricts the emergence of resistance

We next asked whether NAC-mediated enhancement of moxifloxacin lethality occurs *in vivo*. Recent work by Vilcheze *et. al.* (Vilchèze and Jacobs, 2021) directed our choice of murine models: they found little enhancement by NAC on the overall lethality of a combination of moxifloxacin and ethionamide when measured after long incubation using an acute model of murine tuberculosis. Since ROS accelerate killing *in vitro* without increasing the extent of killing (Liu et al., 2012), we shortened the drug treatment time. For this pilot test, we infected BALB/c mice with the clinical MDR strain, *M. tuberculosis* NHN1664. Three weeks after aerosol-mediated infection at a low dose (∼100 bacilli), mice were treated once daily for ten days with a low dose of moxifloxacin (50 mg/kg body weight), NAC (500 mg/kg body weight), or moxifloxacin plus NAC at the same doses as for mono treatments (Fig 8A). After 10 days, moxifloxacin monotherapy reduced the bacillary load by 3-fold and 100-fold in lung and spleen, respectively; NAC alone exhibited no anti-bacterial activity (Fig 8B, 8C). The combination of moxifloxacin plus NAC reduced bacterial burden by another 4- and 12-fold beyond that observed for moxifloxacin alone for lung and spleen, respectively (Fig 8B, 8C). Thus, this preliminary finding indicates that NAC stimulates the lethal action of moxifloxacin *in vivo*.

**Fig 8.**
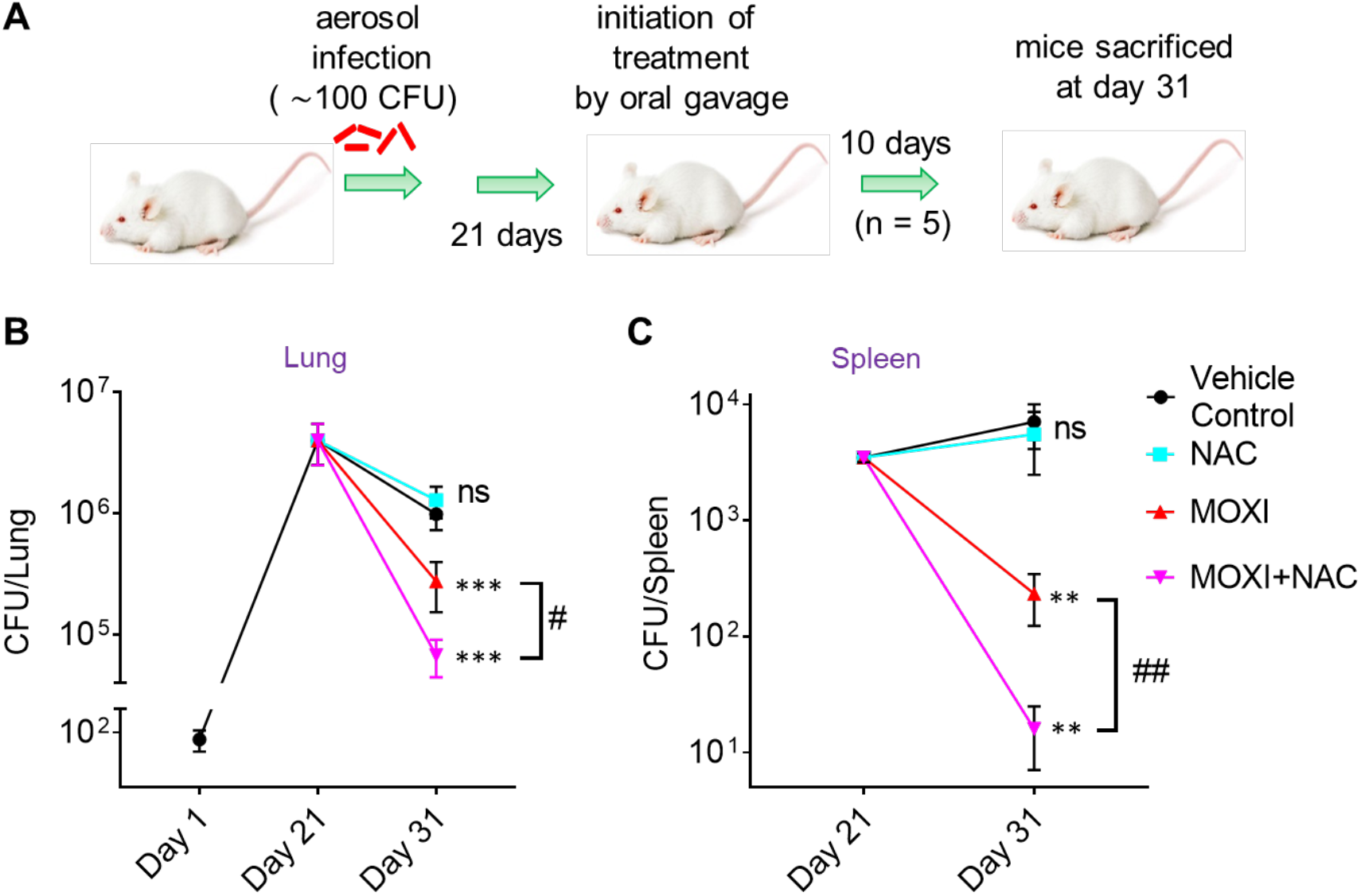
NAC decreases MDR *M. tuberculosis* survival in mice when combined with moxifloxacin. **(A)** Experimental protocol. **(B** and **C)** Bacterial CFUs were enumerated from lungs and spleen at the indicated times. *p* was determined by unpaired two-tailed student’s t-test analyzed relative to vehicle control treatment (** *p* ≤ 0.01, *** *p* ≤ 0.001; ns indicates not significant). Statistical significance between moxifloxacin (MOXI) alone and MOXI + NAC treatment is also shown (# *p* < 0.05, ## *p* ≤ 0.01). Error bars represent standard deviation from the mean of bacterial burden in 5 mice per group.

As a proof of concept. we also measured the effect of NAC on the selection of moxifloxacin-resistant mutants in mice as described (Almeida et al., 2007; Ginsburg et al., 2005). We implanted ∼ 4x 10^3^ *M. tuberculosis* NHN1664 in the lungs of BALB/c mice, and at day 14 post-infection, when the mean lung log_10_ CFU reached ∼ 10^7^, mice received moxifloxacin, NAC, or moxifloxacin plus NAC. After 14 days of post-treatment, 66% of moxifloxacin-treated mice harbored resistant bacteria as compared to 17% when NAC was also present. The combination of moxifloxacin plus NAC brought the number of mutants down to the level seen with untreated animals (Table S7). Enumeration of total resistant colonies confirmed that while moxifloxacin treatment increases emergence of resistance, NAC supplementaion reduces the recovery of moxifloxacin-resistant mutants by 8-fold (Table S7) in our pilot experiment.

## Discussion

The experiments described above show that *M. tuberculosis* responds to moxifloxacin through a lethal stress response characterized by suppressed respiration, elevated NADH levels, and ROS accumulation (shown schematically in Fig 9). Although ROS accumulation is also observed with *E. coli* during lethal stress, early steps in the *M*. *tuberculosis* stress response are distinct. *M. tuberculosis* enters a metabolically quiescent state characterized by increased NADH levels and NADH/NAD^+^ ratio plus reduced respiratory and glycolytic rates. This response is opposite to that seen with *E. coli* (Dwyer et al., 2014; Kohanski et al., 2007; Lobritz et al., 2015). Decelerated respiration may be an adaptive strategy against ROS-inducing agents produced by the host. Production of ROS likely derives from increased NADH/NAD^+^ ratio, which increases the labile, reduced form of Fe to fuel the Fenton reaction. Interfering with the NADH/NAD^+^ increase dampened reductive stress and moxifloxacin lethality, suggesting that NADH disposal pathways could be targeted to enhance fluoroquinolone lethality with *M. tuberculosis*.

**Fig 9.**
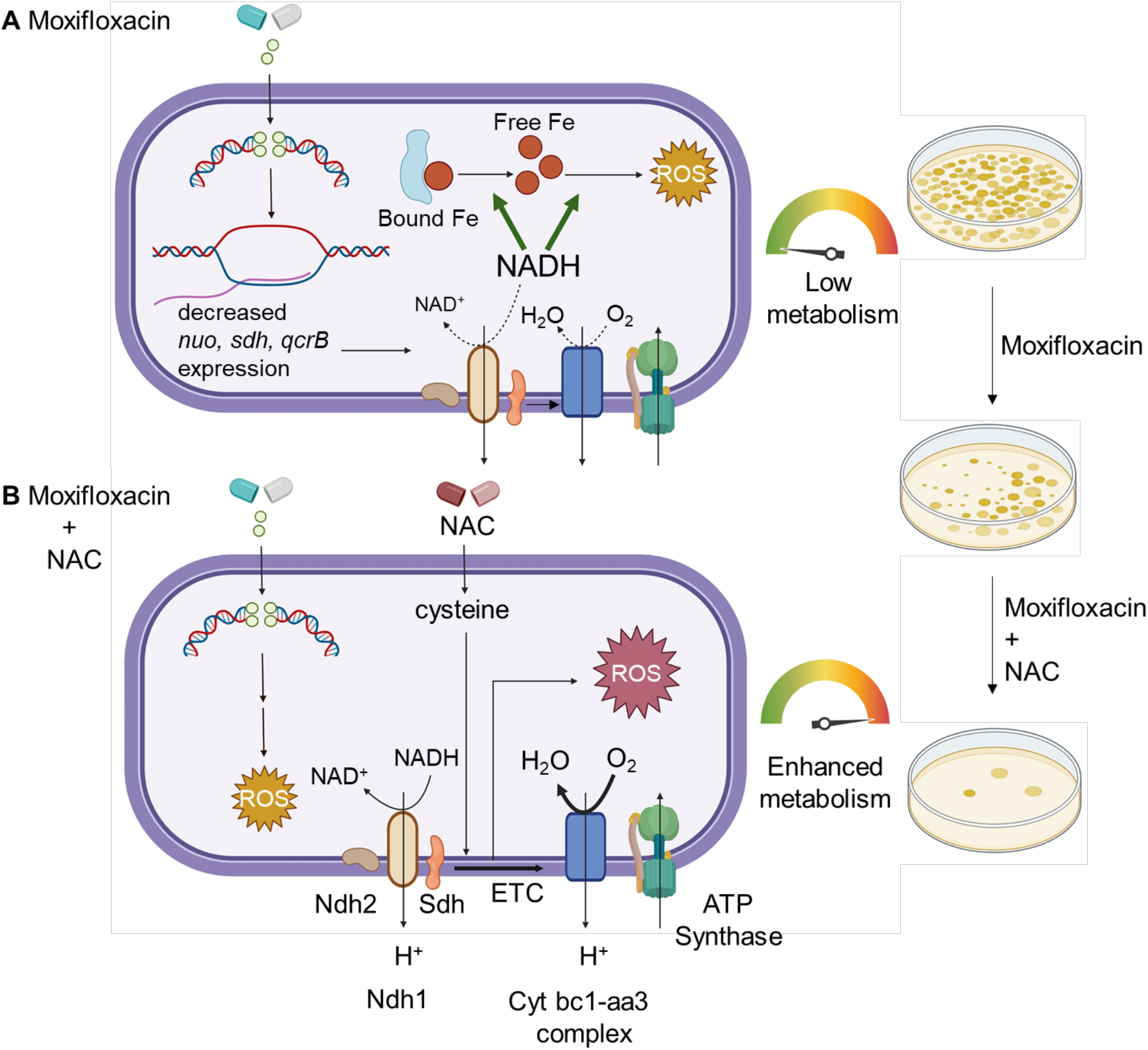
Moxifloxacin-mediated killing of *M. tuberculosis* involves accumulation of NADH-dependent ROS, which is further enhanced by NAC. (a) Moxifloxacin enters *M. tuberculosis* and traps gyrase on DNA as reversible, bacteriostatic drug-enzyme-DNA complexes in which the DNA is broken. The bacterium responds by down-regulating the expression of genes involved in respiration. The transcriptional changes result in reduced rate of respiration. NADH levels and the ratio of NADH to NAD^+^ increase. NADH increases the free Fe^2+^ pool by releasing Fe from ferritin-bound forms and keeps it in a reduced state. ROS damage macromolecules in a self-amplifying process, as indicated by exogenous catalase blocking killing when added after removal of moxifloxacin. (b) Addition of N-acetyl cysteine to cells stimulates respiration and provides more ROS from moxifloxacin-mediated lesions. NAC alone does not induce ROS or trigger death. The additional ROS increases killing by moxifloxacin. Repair of moxifloxacin-mediated lesions, NADH dissipation, Fe sequestration, and ROS detoxification mechanisms contribute to survival.

### ROS contribution to lethality

Studies with *E. coli* indicate that ROS kill bacteria by damaging macromolecules *i.e.* DNA breakage, protein carbonylation, and lipid peroxidation (Drlica and Zhao, 2020). Our data support the general principle that ROS contribute to stress-mediated death of moxifloxacin-treated *M. tuberculosis*: 1) biosensors show an increase in ROS at lethal drug concentrations, 2) iron concentrations increase with moxifloxcin treatment, and 3) two ROS-mitigating agents, thiourea and bipyridyl, reduce oxidative stress and killing. However, the striking increase in survival for *E. coli* at very high quinolone concentrations is associated with a drop in ROS, while this is not the case with *M. tuberculosis*; this difference remains unexplained.

Another difference is the lowering of residual fluoroquinolone lethality by nitrate with hypoxic *M. tuberculosis*: the opposite is observed with norfloxacin-treated *E. coli*. Previous *M. tuberculosis* work has shown that nitrate protects hypoxic bacilli from killing caused by exposure to reactive nitrogen intermediates (Tan et al., 2010), acid stress (Tan et al., 2010), sudden anaerobiosis (Sohaskey, 2008), or inhibition of type II NADH-dehydrogenase (*ndh-2*) by thioridazine (Sohaskey, 2008). Two phenomena may contribute to nitrate-mediated protection. 1) *M. tuberculosis* reduces nitrate to nitrite (Cunningham-Bussel et al., 2013), which oxidizes ferrous Fe to the ferric state, thereby displacing Fe from critical Fe-sulfur clusters and inducing bacteriostasis (Cunningham-Bussel et al., 2013). Nitrite also remodels the *M. tuberculosis* transcriptome similar to several host-imposed stresses that suppress antibiotic-mediated lethality (Cunningham-Bussel et al., 2013) and ensure bacterial survival. 2) Nitrate, as a terminal electron acceptor, recycles or disposes of excess reducing equivalents that accumulate during anaerobic conditions (Shapleigh, 2009). Discovering strategies to halt such adaptive remodeling (*e.g.,* nitrate reduction) in *M. tuberculosis* could serve as adjunct therapy for moxifloxacin with hypoxic bacilli.

### Reductive stress as a source of ROS

When we dissipated NADH-reductive stress by overexpression of *LbNox*, the ROS surge was mitigated, and moxifloxacin lethality was lowered. Production of NADH can also be lowered by stimulating the glyoxylate shunt via isocitrate lyase (*icl*) *−* deletion of *icl* increases the lethal action of isoniazid, rifampicin, and streptomycin (Nandakumar et al., 2014), suggesting that NADH-mediated ROS accumulation may also contribute to the lethal action of these agents. The observed increase in NADPH/NADP^+^ ratio in response to moxifloxacin exposure could assist *M. tuberculosis* in mitigating oxidative stress by activating NADPH-dependent antioxidant systems, thioredoxins, and mycothiol. As shown by microarray data, the increase in NADPH/NADP^+^ ratio could also be a consequence of down-regulation of NADPH-requiring anabolic pathways, such as polyketide/lipid biogenesis, amino acid anabolism, and *de novo* nucleotide synthesis.

In summary, *M. tuberculosis* appears to tolerate lethal stressors of the host immune system by decelerating respiration coupled with dissipation of NADH-reductive stress. The need to dissipate reductive stress may be of general importance, as this type of stress is also generated and amplified by hypoxia, inhibition of aerobic respiration by NO, and catabolism of fatty acids (Singh et al., 2009; Voskuil et al., 2003). *M. tuberculosis* dissipates reductive stress through WhiB3-mediated anabolism of polyketide lipids (Singh et al., 2009). Other bacterial species also dispose of excess reductant, which they do by 1) using nitrate as a terminal electron acceptor during hypoxia (Tan et al., 2010), 2) fueling fermentation under nitrosative stress (Richardson et al., 2008), and 3) secreting phenazines (redox-active polyketides) that generate NAD^+^ from NADH (Price-Whelan et al., 2007). Moxifloxacin is the first anti-mycobacterial agent shown to stimulate NADH-reductive stress.

### Lethality enhancement by N-acetyl cysteine

Vilchèze and Jacobs (Vilchèze and Jacobs, 2021) reported that NAC boosts the lethality of anti-TB drug combinations *in vitro*. We emphasize that a general, enhancing effect on drug-target interactions by NAC is unlikely, because such effects are usually detected by MIC measurement (Drlica and Zhao, 2020): NAC has no effect on moxifloxacin MIC. This observation reinforces the concept that drug-target interactions are mechanistically distinct from downstream ROS-based killing (protection from killing, which is called tolerance when <u>no</u> change in MIC occurs, is distinct from resistance, which is measured as an increase in MIC).

To simplify interpretation of killing data, we focused on a single drug, moxifloxacin, rather than drug combinations. Previous demonstration that NAC stimulates respiration (Vilchèze and Jacobs, 2021) made it a useful tool for examining killing mechanism. NAC increased moxifloxacin-mediated ROS accumulation and lethality with cultured *M. tuberculosis*. Increased killing was initially surprising, because NAC has anti-oxidant properties expected to reduce moxifloxacin lethality (Ezeriņa et al., 2018). It is likely that the effect of NAC depends on the level of stress, since with other bacterial species, genes (*mazF, cpx, lepA*) and treatments (chemical generation of superoxide, H_2_O_2_) are known to be protective at low levels of stress but destructive at high ones (Dorsey-Oresto et al., 2013; Li et al., 2014; Mosel et al., 2013; Rodríguez-Rojas et al., 2020). Indeed, treatment with a sub-lethal concentration of H_2_O_2_ activates the OxyR-regulon in *E. coli* and “primes” the bacterium to conteract subsequent oxidative-stress challenge. At higher concentrations, H_2_O_2_ is cidal (Imlay and Linn, 1986). With the high level of stress produced by moxifloxacin, the respiration-stimulating activity of NAC dominates and adds to ROS derived from NADH-reductive stress.

As expected from *in vitro* studies, NAC increased moxifloxacin lethality with infected macrophages and mice. The kinetic effects of ROS explain why our short-term treatment model with infected mice showed stimulation by NAC, while the longer treatment used previously with antimicrobial combinations did not (Vilchèze and Jacobs, 2021).

We addressed the potential of NAC to suppress the emergence of resistance by measuring its effect on MPC. Although MPC is a bacteriostatic parameter (MIC of the least susceptible mutant subpopulation), highly lethal agents lower it in an animal model of infection, presumably by killing mutants (Cui et al., 2006). Thus, we were not surprised that stimulation of moxifloxacin lethality by NAC lowered MPC. In a murine infection model, NAC reduced the recovery of resistant mutants by 8-fold.

Although MPC for moxifloxacin is lower than concentrations achieved in serum by approved doses (Dong et al., 2000), the clinical situation is likely complex: moxifloxacin shows poor penetration into caseous regions of tubercular granulomas in a rabbit model of experimental tuberculosis (Prideaux et al., 2015; Sarathy et al., 2019). Low, local moxifloxacin concentrations may promote the emergence of fluoroquinolone resistance (Davies Forsman et al., 2020). Thus, NAC supplementation could increase the probability that moxifloxacin concentration is above MPC in vivo, but additional work is required to test this idea.

NAC is a potential candidate as a lethality enhancer, because it is relatively safe for humans (it has been used to alleviate drug-induced toxicity in TB-patients (Baniasadi et al., 2010; Kranzer et al., 2015)). Furthermore, NAC by itself exhibits anti-mycobacterial activity inside macrophages, mice, and guinea pigs (Amaral et al., 2016; Palanisamy et al., 2011), and a small clinical trial in which NAC was administered to TB patients improved host health, immunological response, and early sputum conversion (Mahakalkar et al., 2017; Nagrale et al., 2013). The present work provides a mechanistic foundation for refining NAC-based enhancement of anti-tuberculosis agents. Assessing the general clinical significance of ROS enhancement is also complex, because ROS accelerates killing without lowering bacterial survival at long incubations. This phenomenon has been observed with *S. aureus* (Liu et al., 2012), *E. coli* (Hong et al., 2019), and *M. tuberculosis* (Fig. 2F; S4B). With *M. tuberculosis*-infected mice, NAC stimulates reduction of bacterial load at short but not long incubation times (Vilchèze and Jacobs, 2021). Thus, an appropriate dosing interval must be determined, initially guided by *in vitro* pharmacodynamic studies (Almeida et al., 2007; Firsov et al., 2003; Ginsburg et al., 2005; Gumbo et al., 2004). Relevant tissue concentrations of NAC and moxifloxacin for various doses over time must also be determined before clinical relevance can be assessed. The present report, plus that of Vilchèze and Jacobs (Vilchèze and Jacobs, 2021), encourage such followup work.

## Materials and Methods

### Bacterial strains and culture conditions

*M. tuberculosis* H37Rv and drug-resistant clinical isolates JAL 1934, JAL 2287, JAL 2261, and BND 320 were kind gifts from Dr. Kanury V.S. Rao (International Centre for Genetic Engineering and Biotechnology, Delhi, India). MDR strain NHN1664, originally isolated in China, was obtained from BEI Resources, NIAID, NIH. Strain *Mtb*-roGFP2 was generated by transforming *M. tuberculosis* strain H37Rv with an *E. coli*-mycobacterial shuttle vector, pMV762 (Singh et al., 2006; Steyn et al., 2003). Plasmid pMV762 contains the Mrx1-roGFP2 biosensor construct under control of the *M. tuberculosis hsp60* promoter and a hygromycin-resistance gene as a selection marker. Hygromycin (MP Biomedical, cat. No. 0219417091, Santa Ana, CA) was used at a final concentration of 50 µg/mL. Plasmid (pGMCgS-0×-Ptb38-LbNOX-FLAG-SD1) containing a *lbnox* construct was kind gift from Dr. Dirk Schnappinger and Dr. Sabine Ehrt (Weill Cornell Medicine, New York, USA). Plasmid was electroporated into wild-type *M. tuberculosis* H37Rv to create the *Mtb-LbNox* strain. Streptomycin was used at a final concentration of 25 µg/mL. Plasmid pMV762-Peredox-mcherry was a kind gift from Dr. Ashwani Kumar (Council of Scientific and Industrial Research, Institute of Microbial Technology, Chandigarh, India).

All strains, including laboratory *M. tuberculosis* strain H37Rv and *M. tuberculosis* expressing the Mrx1-roGFP2 redox biosensor, were grown in 7H9 broth supplemented with 0.2% glycerol, 0.05% Tween-80, and ADS (0.5% albumin, 0.2% dextrose, and 0.085% NaCl) with shaking at 180 RPM in a rotary shaker incubator (Lab Therm LT-X; Kuhner, Basel, Switzerland) or on 7H11 agar supplemented with ADS or OADC (ADS plus 0.05% oleic acid and 0.004% catalase (Sigma Aldrich, cat. No. C9322, St Louis, MO) or ADC (ADS plus 0.004% catalase) at 37°C. Bacterial strains expressing Mrx1-roGFP2 were grown in media containing hygromycin.

### Determination of minimal inhibitory concentration (MIC) under aerobic growth conditions

MIC was determined by a resazurin microtiter assay (REMA) using 96-well flat-bottom plates (Padiadpu et al., 2016). *M. tuberculosis* strains were cultured in 7H9+ADS medium (Bhaskar et al., 2014) and grown to exponential phase (OD_600_ = 0.4 to 0.8). Approximately 1×10^5^ bacteria per well were added in a total volume of 200 µL of 7H9+ADS medium. Wells lacking *M. tuberculosis* served as controls. Additional controls consisted of wells containing cells without drug treatment (growth control). After 5 days of incubation at 37°C in the presence of fluoroquinolone (or NAC), 30 µL of 0.02% resazurin (Sigma-Aldrich, cat. No. R7017, St Louis, MO) was added, and plates were incubated for an additional 24 h. Fluorescence intensity was measured using a SpectraMax M3 plate reader (Molecular Devices, San Jose, CA) in the bottom-reading mode with excitation at 530 nm and emission at 590 nm. Percent inhibition was calculated from the relative fluorescence units compared with an untreated control culture; MIC was taken as the lowest drug concentration that resulted in at least 90% reduction in fluorescence compared to the untreated growth control.

### Determination of bacterial survival

To determine the number of viable bacilli, aliquots were removed from cultures, cells were concentrated by centrifugation (4200 *g*, 5 min) to remove drug or treatment compounds, and they were resuspended in an equal volume of medium. Ten-fold dilutions were prepared, and 20 µL aliquots were spotted on drug-free 7H11 agar plates containing 10% ADC or ADS. Plates were incubated for 3-4 weeks at 37°C for CFU enumeration by visual inspection.

Concentration-kill curves were obtained by treating exponentially growing cultures of *M. tuberculosis* (OD_600_ of 0.3 or ≈ 5×10^7^ cells/mL) with various concentrations of moxifloxacin. Tubes were incubated with shaking at 180 RPM for 10 days at 37°C. Aliquots were taken at various intervals, serially diluted, and plated on drug-free agar for CFU enumeration.

LD_90_ (lethal dose) for moxifloxacin under *in vitro,* aerobic-culture conditions was measured by treating 10-mL cultures in 50-mL centrifuge tubes under aerobic conditions (shaking at 180 RPM, 37°C), followed by CFU measurement. A 90% reduction in CFU on treatment day 5, compared with the input control, was defined as LD_90_.

To determine the effect of an Fe-chelator on moxifloxacin-mediated lethality, 250 µM (a non-inhibitory concentration) of bipyridyl (Sigma-Aldrich, cat.no. D216305, St Louis, MO) was added to bacterial cultures 15 min prior to moxifloxacin addition; bipyridyl was present throughout the experiment (i.e., 10 days).

For experiments with thiourea or N-acetyl cysteine (NAC), thiourea (10 mM) or NAC (1 mM) was added 1 h prior to moxifloxacin and was maintained for 2 days or 10 days. Aliquots were taken at indicated times, serially diluted, and plated for CFU enumeration on drug-free agar.

### Post-stressor bactericidal activity of moxifloxacin

*M. tuberculosis* cultures were treated with moxifloxacin for 48 h, the drug was removed by washing, and cells were plated on drug-free 7H11 agar with or without 1 mM NAC or with 2.5% bovine serum albumen (BSA), followed by visual CFU determination.

### Measurement of *E_MSH_* using the Mrx1-roGFP2 redox biosensor

For intra-mycobacterial *E_MSH_* determination during *in-vitro* growth or during *ex-vivo* macrophage-infection conditions, strain *Mtb*-roGFP2 was grown in 7H9 broth or in THP-1 macrophages, respectively. Cultures were then treated with moxifloxacin, and at various times they were treated with 10 mM N-ethylmaleimide (NEM; Sigma-Aldrich, cat. No.-E3876, St Louis, MO) for 5 min at room temperature followed by fixation with 4% paraformaldehyde (PFA; Himedia, cat. No. GRM3660, Mumbai, India) for 1 h at room temperature. Bacilli or infected macrophages were analyzed using a FACSVerse Flow cytometer (BD Biosciences, San Jose, CA). Intramycobacterial *E_MSH_* was determined using the Nernst Equation as described previously (Bhaskar et al., 2014). *E_MSH_* is defined as the standard reduction potential of the MSH/MSSM redox couple (reduced mycothiol to oxidized mycothiol.

The biosensor response was measured by analyzing the fluorescence ratio at a fixed emission (510 nm) after excitation at 405 and 488 nm. Data were analyzed using the BD FACSuite software. These ratiometric data were normalized to measurements with cells treated with 10 mM cumene hydroperoxide (Sigma-Aldrich-cat. No. 247502, St Louis, MO), which reports maximal oxidation of the biosensor, and 20 mM dithiothreitol (Sigma-Aldrich, cat. No. D9779, St Louis, MO), which reports maximal reduction. 10,000 events per sample were analyzed.

### ROS measurement using CellROX Deep Red

Cultures of exponentially growing *M*. *tuberculosis* at an initial OD_600_ of 0.3 were treated with various concentrations of moxifloxacin for 48 h. Sterile 50 mL-capacity polypropylene centrifuge tubes (Abdos, cat no. P10404, Roorkee, India), containing 10 mL of culture, were incubated in a shaker incubator (180 RPM, 37°C). Then cells were harvested by centrifugation (4200 *g* for 5 min) and resuspended in 100 µL of growth medium. As per manufacturer’s instructions, CellROX Deep Red (Invitrogen, cat. No. C10422, Waltham, MA) was added to a final concentration of 5 µM, and cells were agitated on a rocker (Biobee Tech, Bangalore, India) for 30 min. After incubation, cells were washed to remove residual dye by centrifugation (4200 *g* for 5 min). Cells were resuspended in 300 uL phosphate-buffered saline, pH 7.4) and then fixed by addition of 4% paraformaldehyde (PFA) for 1 h at room temperature. Fluorescence was measured at a fixed emission (670 nm) after excitation with a red laser (640 nm) using a BD FACSVerse Flow cytometer (BD Biosciences, San Jose, CA) with 10,000 events per sample. No autofluorescence was observed.

### Determination of DNA damage by TUNEL assay

DNA damage was measured by using In Situ Cell Death Detection Kit, TMR red (Roche Molecular Biochemicals, Indianapolis, IN, Cat. No.-12156792910), which is based on TUNEL (TdT-mediated dUTP-X nick end labeling) assay (Fan et al., 2018; Vilchèze et al., 2013). An equal number of cells (based on OD_600_) were taken at various times of treatment with moxifloxacin, washed once by centrifugation, and fixed in 2% paraformaldehyde (PFA). PFA was removed by washing cells, followed by resuspension in 2% sodium dodecyl sulfate (SDS) and a second wash by centrifugation. DNA double-strand breaks were labeled in 100 μl of TUNEL reaction mix for 3-4 h. Cells incubated with label solution only (no terminal transferase) were used as negative controls. Fluorescence was measured at a fixed emission (585 nm) after excitation with green-yellow laser (561 nm) using a BD FACSAria Flow cytometer (BD Biosciences, San Jose, CA). 10,000 events were acquired per sample.

### Lipid Peroxidation Assay

Cultures were grown to mid-log phase (OD_600_= 0.6-0.8). Lipid hydroperoxides were quantified from cell pellets after centrifugation (4000 *g,* 5 min), using FOX2 reagent (Nambi et al., 2015). Briefly, cell pellets were resuspended in 1:2 chloroform/methanol and mixed by vortexing. Next, chloroform and water were added at a 1:1 mixture The samples were then centrifuged to separate the water and organic phases. The organic phase was collected and washed twice with water. 200 µl of the organic phase was incubated with 1 ml of FOX2 reagent in the dark for 1 h at 22°C. Lipid hydroperoxides were measured spectrophotometrically at 560 nm and normalized to culture turbidity (OD_600_).

### Macrophage preparation and infection by *M. tuberculosis*

The human monocytic cell line THP-1 was grown in RPMI1640 medium (Cell Clone, Manassas, VA) supplemented with 10% heat-treated (55°C) fetal bovine serum (MP Biomedical, cat. no. 092916754, Santa Ana, CA). A total of 3×10^5^ cells/well was seeded into a 24-well cell-culture plate. THP-1 monocytes were stimulated to differentiate into macrophage-like cells by treatment with 20 ng/mL phorbol 12-myristate 13-acetate (PMA; Sigma-Aldrich Co., St Louis, MO) for 18-20 h at 37°C and incubated further for 48 h to allow differentiation (Bhaskar et al., 2014). The resulting macrophage-like cells were infected with *Mtb*-roGFP2 or MDR *M. tuberculosis* NHN1664 expressing Mrx1-roGFP2 at a multiplicity of infection (MOI) of 10 and incubated for 4 h at 37 °C in 5% CO_2_. After Infection, extracellular bacteria were removed by washing three times (by centrifugation) with phosphate-buffered saline (PBS; 137 mM NaCl, 2.7 mM KCl, 10 mM Na_2_HPO_4_, and 1.4 mM KH_2_PO_4_, pH 7.4). Moxifloxacin (Sigma-Aldrich, cat. No. PHR1542, St Louis, MO), with or without NAC (Sigma-Aldrich, cat. No. A7250, St Louis, MO), was added to infected cells that were incubated for various times. For CFU determination, infected cells were lysed in 7H9 medium containing 0.06% sodium dodecyl sulfate (SDS); dilutions were prepared using 7H9 medium, and aliquots were plated on 7H11+OADC agar plates (Bhaskar et al., 2014). Plates were incubated at 37 °C for 3-4 weeks before colonies were counted.

### Nutrient starvation

Starvation was achieved as previously described (Gengenbacher et al., 2010). Briefly, cultures of *M. tuberculosis* H37Rv were grown to exponential phase in roller-culture bottles (Corning, cat no.-430518, Corning, NY) containing Middlebrook 7H9 medium, supplemented with ADS and 0.05 % Tween 80, at 37°C with rolling at 6 RPM in a roller incubator (120 Vac Roll-In Incubator; Wheaton, Millville, NJ). Cultures grown to OD_600_ ≈ 0.2 were harvested by centrifugation (4000 *g* for 5 min) followed by two washes with phosphate-buffered saline (PBS; pH 7.4) supplemented with 0.025 % Tween 80 (PBST). Bacterial cells were diluted to a final OD_600_ of 0.1 in PBS. 50 mL of this suspension was transferred into a roller-culture bottle and incubated for 14 days to achieve starvation conditions.

### Hypoxia

For determination of moxifloxacin lethality under hypoxic conditions, bacterial cultures (OD_600_ = 0.1) were placed in Vacutainer tubes (Becton Dickinson, cat. no. 367812, Franklin Lakes, NJ) followed by incubation for 14-21 days at 37°C (Taneja and Tyagi, 2007). A high cell density (OD_600_ = 0.1) was used for rapid achievement of hypoxia, which was observed as decolorization of methylene blue (final concentration was 1.5 μg/mL) in the culture medium. When hypoxia was established, moxifloxacin was added to cultures anaerobically. Metronidazole at 10 mM and isoniazid at 10 µM were used as positive and negative controls, respectively. Drugs were injected in volumes of 100 µL in phosphate-buffered saline following passage of argon through the drug solution to remove residual oxygen. Hypoxic cultures were treated with drugs for five days, similar to the incubation time for MIC determination with aerobically growing cells. After treatment, Vacutainer tubes were unsealed, and end-point bacterial survival was determined by plating on drug-free agar, incubating for 3-4 weeks at 37°C, and visually enumerating colonies.

### Effect of nitrate on survival of hypoxic bacteria

Survival during nitrate-dependent respiration was achieved by supplementing *M. tuberculosis* cultures with 5 mM sodium nitrate. Nitrate was added when cultures were placed in Vacutainer tubes before hypoxia. All other conditions were as described above for hypoxia.

### Microarray data analysis

Gene expression data from GSE71200 (Gene Expression Omnibus, NCBI or GEO) was used for analyzing the response of *M. tuberculosis* to moxifloxacin (Ma et al., 2015). The data were expressed as a two-channel microarray with control and drug-exposed *M. tuberculosis* stained with Cy3 and Cy5 dyes, respectively. We used GSM1829746, GSM1829747, and GSM1829748, which consist of published data for *M. tuberculosis* exposed to 2X, 4X, or 8X MIC (1X MIC = 0.4 µM) of moxifloxacin for 16 h. Normalized expression data for GSE71200 was downloaded from GEO (Barrett et al., 2013; Edgar et al., 2002), and probe IDs were mapped to the respective gene-IDs. The expression levels for genes having multiple probes were averaged, and genes lacking data were removed from further analysis. Differentially expressed genes (DEGs) were defined as genes that were upregulated or downregulated by at least 2-fold in all 3 moxifloxacin treatment conditions.

For overlap analysis of differentially expressed genes in moxifloxacin-exposed bacteria compared with other stress-induced conditions (log_2_ fold change >1 or <-1; *p* < 0.01 are considered differentially expressed genes in stress conditions), GeneOverlap (v1.22.0) package from R (v3.6.3) was used (Core-Team, 2018; Li and Sinai, 2019). It uses Fisher’s exact test to find statistical significance by calculating the *p*-value and the odds ratio for the overlap (*p*-value < 0.05 and an odds ratio of > 1 were taken as the significance thresholds).

### OCR and ECAR measurements

To measure basal oxygen consumption rate (OCR) and extracellular acidification rate (ECAR), log-phase *M. tuberculosis* cultures (OD_600_= 0.6-0.8) were briefly (one day) incubated in 7H9 medium containing the non-metabolizable detergent tyloxapol (MP Biomedical, cat. No.157162, Santa Ana, CA) and lacking ADS or a carbon source. These cultures were then passed 10 times through a 26-gauge syringe needle followed by centrifugation at 100 *g* for 1-2 mins to remove clumps of bacterial cells. The resulting single-cell suspensions of bacteria at 2×10^6^ cells/well were placed in the bottom of wells of a Cell-Tak (Corning, cat. No. 354240, Corning, NY)-coated XF culture plate (Agilent/Seahorse Biosciences, Santa Clara, CA). Measurements were performed using a Seahorse XFp analyzer (Agilent/Seahorse Biosciences, Santa Clara, CA) with cells in unbuffered 7H9 growth medium (pH 7.35 lacking monopotassium phosphate and disodium phosphate) containing glucose (2 mg/mL) as a carbon source. OCR and ECAR measurements were recorded for ∼21 min (3 initial baseline readings) before addition of moxifloxacin (1X, 2.5X or 10X MIC; 1X MIC = 0.5 µM), which was delivered automatically through the drug ports of the sensor cartridge (Wave Software, Agilent Technologies, Santa Clara, CA). NAC or CCCP was similarly added through drug ports at times indicated in figures. OCR and ECAR were measured for an additional 6 h in the absence or presence of moxifloxacin and/or NAC. Changes in OCR and ECAR readings triggered by moxifloxacin were calculated as a percentage of the third baseline reading for OCR and ECAR taken prior to drug injection.

### ROS measurement under Fe-depletion conditions

Iron starvation was as described previously (Kurthkoti et al., 2017). Briefly, log-phase *M. tuberculosis* cultures were grown in minimal medium (0.5% (wt/vol) asparagine, 0.5% (wt/vol) KH_2_PO4, 2% glycerol, 0.05% Tween 80, 10% ADS, 0.5 mg/L of sterile ZnCl_2_, 0.1 mg/ L of MnSO_4_, and 40 mg/L of MgSO_4_) to early stationary phase (OD_600_ of ∼1). To remove metal ions, the minimal medium was treated with 5% Chelex-100 (Bio-Rad, cat. No. 142-2842, Hercules, CA) for 24 h with gentle agitation. This Fe-depleted culture was diluted further to OD_600_ of 0.1 in the same medium and allowed to grow to early stationary phase to deplete stored Fe. The Fe-depleted cells were further treated with 50 µg/ml of freshly prepared deferoxamine mesylate (Sigma Aldrich, cat. No. D9533, St Louis, MO)-containing minimal medium for 6 days. The cells were washed and diluted in minimal medium containing 80 µM FeCl_3_ or 80 µM FeCl_3_ + 10 mM thiourea for an additional 4 days. ROS was quantified by CellROX Deep Red dye using Flow cytometry as described in the section titled ROS measurement using CellROX Deep Red.

### Cellular Fe Estimation

Cell-free Fe levels were measured using the ferrozine-based assay as described previously (Fish, 1988; Vilchèze et al., 2017). Briefly, *M. tuberculosis* cultures were grown to an OD_600_ of 0.3-0.4 followed by treatment with 0X, 1X, 2.5X, or 10X MIC of moxifloxacin. After 48 h of treatment, cells were harvested and washed twice with ice-cold PBS. The cell pellets were resuspended in 1 mL of 50 mM NaOH and lysed using a bead beater. The cell lysate sample (300 µl) was mixed with 10 mM HCl (300 µL) followed by addition of Fe-detection reagent (6.5 mM Ferrozine, 6.5 mM Neocuproine, 1 M ascorbic acid, and 2.5 M ammonium acetate in water) (90 µl). The reaction mix was incubated at 37°C for 30 min, and then absorbance at 562 nm was measured. The cellular Fe concentration was determined by plotting the absorbance values against a standard curve of FeCl_3_ concentration gradient and normalized to protein content. Protein concentration was determined using the Pierce BCA Protein Assay Kit (Thermo Scientific, cat. no. 23225, Rockford, IL).

### Estimation of NAD^+^, NADP^+^, NADH, and NADPH

*M. tuberculosis* H37Rv was grown to OD_600_ ∼0.35 and treated with 1X, 2.5X or 10X MIC of moxifloxacin for 48 h. Pyridine nucleotide levels were determined by a redox-cycling assay (Chawla et al., 2012; Singh et al., 2009). Protein concentration of NAD^+^ or NADH extracts was determined using the Pierce BCA Protein Assay Kit (Thermo Scientific, cat. no. 23225, Rockford, IL) to normalize NAD(P)H and NAD(P)^+^ concentrations.

### Analysis of mixtures containing moxifloxacin plus and minus NAC by thin layer chromatography (TLC)

Thin-layer chromatography (TLC) was performed using silica gel 60 GF_254_ precoated aluminium backed plates (0.25 mm thickness; Merck, Darmstadt, Germany), and visualization was accomplished by irradiation with UV light at 254 nm. Stock solutions of moxifloxacin (10 mM) and NAC (0.2 M, 2 M and 20 M) were prepared independently in DMSO and phosphate buffer (PB, pH 7.4, 10 mM), respectively. In a typical incubation, moxifloxacin (100 µL at a final concentration of 2 mM) was independently reacted with various concentrations of NAC (5 µL, final concentration 2 mM, 20 mM, or 200 mM) in 395 µL PB (pH 7.4, 10 mM). In a control experiment, moxifloxacin (100 µL, final concentration 2 mM) was added to 400 µL PB (pH 7.4, 10 mM). The mixtures were incubated at 37 °C on an Eppendorf ThermoMixer Comfort (800 rpm). The reactions were monitored by spotting aliquots (5 µL) from the incubation mixtures onto the TLC plate at designated times. The solvent system used was MeOH and CHCl_3_ (1:9).

### Analysis of mixtures containing moxifloxacin plus and minus NAC by NMR

Deuterated phosphate buffer (PB, 10 mM) was prepared by dissolving monobasic potassium phosphate (KH_2_PO_4_, 4 mg) and dibasic potassium phosphate (K_2_HPO_4_, 12 mg) in deuterated water (D_2_O), and the pH was adjusted to 7.4 using 40% (w/w) sodium deuteroxide solution in D_2_O (Sigma-Aldrich, cat. No. 151882, St Louis, MO). Stock solutions of moxifloxacin (10 mM) and NAC (0.2 M) were prepared in DMSO and deuterated phosphate buffer (PB, pH 7.4, 10 mM) respectively. In a typical reaction, moxifloxacin (200 µL, 2 mM final concentration) was incubated with NAC (10 µL, 2 mM final concentration) in 790 µL deuterated PB (pH 7.4, 10 mM). In a control experiment, moxifloxacin (200 µL, 2 mM final concentration) was added to 800 µL deuterated PB (pH 7.4, 10 mM). The incubation mixtures were incubated at 37 °C in an Eppendorf ThermoMixer Comfort (800 rpm). An aliquot (0.5 mL) of the incubation mixture was taken after 1 h of incubation, and ^1^H NMR and ^19^F NMR spectra were recorded. ^1^H NMR was acquired with 64 scans on a Jeol 400 MHz spectrometer using deuterated water (D_2_O; Sigma-Aldrich) as an internal standard. Chemical shifts (δ) were reported in ppm downfield from D_2_O (δ = 4.79 ppm) for ^1^H NMR. ^19^F spectra was recorded on a JEOL (376 MHz) using an external reference (*α*, *α*, *α*-trifluorotoluene, δ_F_ = −63.72 ppm).

### Fluorescence-based detection of intracellular levels of moxifloxacin in wild-type *M. smegmatis*

Stock solutions of moxifloxacin (0.05 mM) and NAC (200 mM) were prepared in DMSO and phosphate buffer (pH 7.4, 10 mM), respectively. Wild-type *M. smegmatis* cultures were grown in Middlebrook 7H9 broth supplemented with glycerol (0.2%) and Tween-80 (0.1%). Exponential-phase cultures of wild-type *M. smegmatis,* grown to OD_600_ of 0.3 (985 μL), were transferred to 1.5 mL Eppendorf tubes and either left untreated or treated with NAC (5 μL, final concentration 1 mM) for 1 h prior to addition of 2x MIC of moxifloxacin (10 μL, final concentration 0.5 μM). Treated cells were incubated with shaking at 180 rpm in a rotary shaker incubator at 37 °C for 48 h. The cell suspension was then centrifuged at 9,391 *g* at 4 °C for 15 min, and the pellet was washed twice with 1x phosphate buffer saline (PBS) and resuspended in 1 mL PBS in a microcentrifuge tube. The cells were lysed by sonication using a 130-watt ultrasonic processor (Vx 130W) by stepping a microtip with a 4 min pulse on-time (with 3s ON and 3s OFF pulse, 60% amplitude, 20 kHz frequency) under ice-cold conditions. An aliquot (100 μL) of whole-cell lysate was withdrawn from the above samples and dispensed into a 96-well microtiter plate. Fluorescence ascribed to moxifloxacin (λ_ex_ = 289 nm and λ_em_ = 488 nm), recovered in NAC-untreated or-pretreated *M. smegmatis* lysates, was recorded using an Ensight Multimode Plate Reader (PerkinElmer, India). Readings were collected from the top with 25 flashes per well and with a focus height adjusted to 9.5 mm.

### Intracellular levels of moxifloxacin in *M. smegmatis* using LC/MS

Stock solutions of moxifloxacin (0.05 mM) and NAC (200 mM) were prepared as described above. Whole-cell lysates of wild-type *M. smegmatis* treated with moxifloxacin (0.5 μM, 2x MIC) plus or minus NAC (1 mM) were prepared as described in the fluorescence-based method employed for the detection of moxifloxacin. An aliquot (100 μL) of whole-cell lysate was withdrawn from the incubation mixtures, diluted with methyl alcohol (100 μL), and centrifuged at 9,390 x *g* at 4 °C for 15 min. The supernatant fluids (50 μL) were removed, diluted with methyl alcohol (50 μL), and analyzed by LC/MS. All measurements were performed using the positive ion mode with high-resolution, multiple reaction monitoring (MRM-HR) analysis with a Sciex X500R quadrupole time-of flight (QTOF) mass spectrometer fitted with an Exion UHPLC system having a Kinetex 2.6 mm hydrophilic interaction liquid chromatography (HILIC) column (100 Å particle size, 150 mm length, and 3 mm internal diameter; Phenomenex, Intek Chromasol Pvt. Ltd., India). Nitrogen was used as the nebulizer gas, with nebulizer pressure set at 50 psi. MS was calibrated in the positive mode, and samples were analyzed with the following parameters: Mode: electrospray ionization (ESI), ion source gas 1 = 40 psi, ion source gas 2 = 50 psi, curtain gas = 30, CAD gas = 7, spray voltage = 5500 V, and temperature = 500 °C. The MRM-HR mass spectrometry parameters were moxifloxacin (Q1, M + H^+^) = 402.18, moxifloxacin-NAC adduct (Q2, M + H^+^) = 565.21, declustering potential = 80 V, collision energy = 20 V, collision exit potential = 5 V, and accumulation time = 0.24 s. The LC runs were for 30 min with a gradient of 100% solvent A (0.1% HCOOH in water) for 5 min, linear gradient of solvent B (acetonitrile 0% to 100%) for 25 min followed by 100% solvent A for 5 min, all at a flow rate of 0.5 mL per min.

### Determination of Mutant Prevention Concentration (MPC)

MPC of moxifloxacin with *M. tuberculosis* NHN1664 was determined by methods described previously (Dong et al., 2000; Singh et al., 2017). Cultures of MDR strain *M. tuberculosis* NHN1664 were grown to OD_600_ 0.6-0.7. Approximately 2.5×10^9^ bacilli were plated on 7H11 agar containing either moxifloxacin (2X, 4X, or 8X MIC) alone or in combination with NAC at either 1 mM or 2 mM. Resistant colonies were enumerated after incubation at 37°C for 8 weeks. The drug-resistant phenotype was confirmed by replating on drug-containing 7H11-agar plates. MPC was determined as the lowest concentration of drug that prevented bacterial colony formation when >2×10^9^ bacteria were plated on drug-containing 7H11 plates. Mutation frequency with moxifloxacin was calculated as the number of mutants appearing on drug-containing plates divided by the viable input bacteria.

### Determination of drug-tolerant *M. tuberculosis* population *ex vivo*

Murine bone-marrow-derived macrophages (BMDMs) were infected with Mrx1-roGFP2-expressing *M. tuberculosis* H37Rv at a multiplicity of infection of 10. After washing off extracellular bacteria, infected macrophages were left untreated for 24 h for the emergence of the drug-tolerant population. At 24 h post-infection, macrophages harbouring *E*_MSH_-reduced and *E*_MSH_-basal *M. tuberculosis* were sorted using BD FACS Aria™ Fusion Flow Cytometer. The sorted, infected BMDM cells were seeded into 96-well plates followed by treatment with 3X MIC of moxifloxacin (1X MIC = 0.5 µM) for 48 h. After treatment, cells were lysed with 0.05% sodium dodecyl sulfate (SDS) in 7H9 medium, serially diluted, and plated on 7H11-OADC agar plates. Plates were incubated at 37°C for 3-4 weeks before colony forming units (CFUs) were enumerated.

### *In vivo* drug efficacy

All animal studies were executed as per guidelines prescribed by the Committee for the Purpose of Control and Supervision of Experiments on Animals, Government of India, with approval from the Institutional Animal Ethical Committee and Biosafety Level-3 Committee. 6- to 8-week-old female pathogen-free BALB/c mice were infected via a low-dose aerosol exposure to the *M. tuberculosis* MDR strain NHN1664 using a Madison chamber aerosol generation instrument. The short-course mouse model of infection was performed as described previously (Lenaerts et al., 2003). At day one post infection, three mice were sacrificed to verify implantation of ∼100 CFU of bacteria per mouse. Mice were randomly divided into 4 groups (n = 5 per group). Feed and water were given *ad libitum*. Treatment with moxifloxacin and/or NAC started 21 days post-infection and continued for 10 days; untreated control mice received water rather than the two test compounds. Treated mice were administered moxifloxacin (50 mg/kg of body weight), NAC (500 mg/ kg of body weight) and a combination of moxifloxacin and NAC at 50 mg/kg and 500 mg/kg body weight, respectively, by oral gavage, once daily. Five infected mice were sacrificed at the start of treatment as pretreatment controls. After treatment, mice were sacrificed, and the lungs and spleen were harvested for measurement of bacterial burden. CFUs were determined by plating appropriate serial dilutions on 7H11 (supplemented with OADC) agar plates and counting visible colonies after 3–4 weeks of incubation at 37°C. Data were normalized to whole organ.

### Detection of moxifloxacin-resistant mutants in murine model of infection

6- to 8-week-old female, pathogen-free BALB/c mice were infected via a high-dose aerosol exposure to the *M. tuberculosis* MDR strain NHN1664 using a Madison chamber aerosol generation instrument. At day one post infection, three mice were sacrificed to determine implantation of >10^3^ CFU of bacteria per mouse. Mice were then randomly divided into 4 treatment groups, with 6 mice each in the vehicle control group, NAC-only group, groups receiving either moxifloxacin or a combination of NAC and moxifloxacin. Feed and water were given *ad libitum*. Treatment with moxifloxacin and/or NAC started 14 days post-infection and continued for 14 days; untreated control mice received water rather than the two test compounds. Treated mice were administered moxifloxacin (50 mg/kg of body weight), NAC (500 mg/ kg of body weight) and a combination of moxifloxacin and NAC at 50 mg/kg and 500 mg/kg body weight, respectively, by oral gavage, once daily. 5 infected mice were sacrificed at the start of treatment as pre-treatment controls. After treatment, mice were sacrificed, and the lungs and spleen were harvested together. Moxifloxacin-resistant mutants were selected by plating the undiluted homogenates (lung+spleen) on five 7H11 plates containing 0.5 μM of moxifloxacin (4X MIC). Plates were incubated for 3–4 weeks at 37°C.

### Statistical analysis

Statistical analysis was performed using GraphPad Prism version 8.4.3 software. Mean and standard deviation values were plotted as indicated in figure legends. A *p-*value of less than 0.05 was considered significant. Statistical significance was determined by unpaired two-tailed student’s t-test; either one-way or two-way ANOVA was performed where comparison of multiple groups was made.

## Acknowledgements

We thank the following for critical comments on the manuscript: Lanbo Shi, Bo Shopsin, and Xilin Zhao.

## Declaration

The funders had no role in study design, data collection and analysis, decision to publish, or preparation of the manuscript. The authors have declared that no competing interests exist.

