## supplemental fig for "Moxifloxacin-mediated killing of *Mycobacterium tuberculosis* involves respiratory downshift, reductive stress, and ROS accumulation"

[#] Equal Contribution

### Contents


#### Supplementary Tables

**Table S1.** Susceptibility of *M. tuberculosis* to fluoroquinolones under *in vitro* aerobic growth conditions.

**Table S2.** Lethality of fluoroquinolones with *M. tuberculosis* under dormancy-inducing conditions.

**Table S3.** Functional categorization of genes deregulated in *M. tuberculosis* by moxifloxacin**.**

**Table S4.** Summary of overlap between moxifloxacin exposure (number of differentially expressed genes = 359) and various stress conditions**.**

**Table S5.** Susceptibility of *M. tuberculosis* strains to diverse anti-TB drugs under *in vitro* aerobic growth conditions.

**Table S6.** Mutant prevention concentration (MPC) of moxifloxacin with NAC against MDR *M. tuberculosis* strain NHN1664.

**Table S7.** N-acetyl cysteine decreases recovery of moxifloxacin-resistant mutants in infected BALB/c mice.

#### Supplementary Figures

**Fig. S1.** Mrx1-roGFP2 measures redox changes in *M. tuberculosis* in response to $H_2O_2$.

**Fig. S2**. Effect of moxifloxacin and NAC on redox biosensor oxidation in *M. tuberculosis* cultures.

**Fig. S3.** Moxifloxacin treatment increases markers of oxidative stress (DNA double strand breaks and lipid hydroperoxides) in *M. tuberculosis*.

**Fig. S4.** Effect of various fluoroquinolones on ROS production in *M. tuberculosis* cultures.

**Fig. S5**. Increased free Fe concentration leads to ROS generation in *M. tuberculosis.*

**Fig. S6.** Moxifloxacin treatment increases free Fe pools in *M. tuberculosis* that are decreased by NADH dissipation.

**Fig. S7.** Reduction in moxifloxacin lethal activity by the ROS scavenger thiourea.

**Fig. S8.** Addition of extra bovine serum albumin to agar plates does not protect *M. tuberculosis* from moxifloxacin-mediated lethality.

**Fig. S9.** Moxifloxacin lethality during nutrient starvation and hypoxia.

**Fig. S10.** Determination of lethal dose ($LD_{90}$) of moxifloxacin with aerobically cultured *M. tuberculosis.*

**Fig. S11.** Heat map showing qRT-PCR validation of 28 randomly selected genes differentially regulated in the microarray data.

**Fig. S12.** Heat-map showing differential gene overlap between moxifloxacin-exposed *M. tuberculosis* and various stress conditions.

**Fig. S13.** Decelerated respiration in *M. tuberculosis* upon moxifloxacin treatment is not due to bacterial death.

**Fig. S14.** Low moxifloxacin concentrations decelerate respiration rate in *M. tuberculosis.*

**Fig. S15.** Ethambutol treatment does not affect respiration rate in *M. tuberculosis*.

**Fig. S16.** Moxifloxacin decreases total NAD/H (NADH+$NAD^+$) in *M. tuberculosis*.

**Fig. S17.** Moxifloxacin increases NADH/NAD+ ratios in *M. tuberculosis* detected by Peredox

**Fig. S18.** $NADP^+$ and NADPH levels in moxifloxacin-treated *M. tuberculosis.*

**Fig. S19.** Total NAD/H (NADH+$NAD^+$) was not affected by overexpression of *LbNox* in *M. tuberculosis*.

**Fig. S20.** Constitutive expression of *LbNox* does not impair metabolism and growth of rate of *M. tuberculosis*.

**Fig. S21.** NAC has no effect on moxifloxacin MIC with *M. tuberculosis*.

**Fig. S22.** Addition of NAC after removal of moxifloxacin increases killing of *M. tuberculosis.*

**Fig. S23**. Several tests argue against formation of adducts between NAC and moxifloxacin (MOXI).

**Fig. S24.** Moxifloxacin-induced oxidative shift in $E_{MSH}$ of *M. tuberculosis* NHN1664 during infection of THP-1 cells.

**Fig. S25.** Intramacrophage $E_{MSH}$-reduced *M. tuberculosis* shows reduced killing by moxifloxacin.

**Dataset S1.** List of deregulated genes in *M. tuberculosis* after treatment with 2X, 4X and 8X MIC of moxifloxacin (1X MIC = 0.4 μM) for 16 h.

#### Supplementary References

#### Supplementary Tables

##### Table S1. Susceptibility of *M. tuberculosis* to fluoroquinolones under *in vitro* aerobic growth conditions

| Strain | Susceptibility characteristic | MIC in µM (µg/mL) | | |
|---|---|---|---|---|
| | | **MOXI[1]** | **LEV[1]** | **CIP[1]** |
| H37Rv | Susceptible | 0.5 (0.21) | 1(0.36) | 2 (0.66) |
| *Mtb*-roGFP2 | Susceptible | 0.5 (0.21) | ND | ND |
| BND-320 | INH[1] monoresistant | 0.5 (0.21) | 1 (0.36) | 2 (0.66) |
| JAL-2287 | MDR[1] | 0.25 (0.1) | 1 (0.36) | 2 (0.66) |
| JAL-1934 | MDR[1] | 0.5 (0.21) | 1 (0.36) | 4 (1.3) |
| JAL-2261 | MDR[1] | 0.5 (0.21) | 1 (0.36) | 4 (1.3) |
| NHN1664 | MDR[1] | 0.125 (0.05) | ND | ND |
| H37Rv-*LbNox* | Susceptible | 0.5 (0.21) | ND | ND |
| Moxi$^R$ | Resistant to MOXI | 1.25 (0.53) | ND | ND |

[1]Abbreviations: MOXI, moxifloxacin; LEV, levofloxacin; CIP, ciprofloxacin; INH, isoniazid; MDR, multidrug resistant; ND: not determined.

##### Table S2. Lethality of fluoroquinolones (FQ) with *M. tuberculosis* in dormancy-inducing conditions

| **FQ** | MIC[1] | MBC[2] | $LCC_{90}$[3] (Starvation) | $WCC_{90}$[4] (Hypoxia) |
|---|---|---|---|---|
| | Concentration in µM | | | |
| **Ciprofloxacin** | 2 | 2 | >32 | ND[5] |
| **Levofloxacin** | 1 | 1 | >32 | ND |
| **Moxifloxacin** | 0.5 | 0.5 | >32 | 10 |

[1]MIC was determined as 90% prevention of resazurin color change compared to untreated control.
[2]MBC was defined as concentration ≥MIC which prevents colonies from appearing when 1/10 volume from MIC plate is regrown on 7H11 agar.
[3]Loebel cidal concentration ($LCC_{90}$): 90% reduction in viable bacilli under nutrient starvation (Gengenbacher et al., 2010).
[4]Wayne cidal concentration ($WCC_{90}$): 90% decrease in viable bacilli under hypoxia (Gengenbacher et al., 2010).
[5]ND: not determined.

**Table S3. Functional categorization of genes deregulated in *M. tuberculosis* by moxifloxacin.**

| Functional category | % genes in each category | |
|---|---|---|
| | Genome | Transcriptome |
| Cell wall and cell processes | 19.1 | 15.9 |
| Conserved hypotheticals | 25.8 | 30.4 |
| Information pathways | 6.02 | 12.9 |
| Intermediary metabolism and respiration | 23.2 | 16.2 |
| Lipid metabolism | 6.74 | 6.68 |
| PE/PPE | 4.16 | 1.95 |
| Regulatory proteins | 4.91 | 3.89 |
| Unknown | 0.37 | 0.27 |
| Virulence, detoxification, adaptation | 5.95 | 5.29 |
| Insertion sequences and phages | 3.64 | 6.26 |

**Table S4: Summary of overlap between moxifloxacin exposure (number of differentially expressed genes = 359) and various stress conditions.**

| Stress condition | No. of differentially expressed genes in the stress condition[2] | Intersection with Moxifloxacin condition | *p*-value of gene-set overlap | Odds Ratio | Reference |
|---|---|---|---|---|---|
| Acidic pH5.5 | 212 | 17 | 0.72 | 0.88 | (Rohde et al., 2007) |
| Hypoxia DosR | 46 | 0 | 1 | 0 | (Rustad et al., 2008) |
| $H_2O_2$ stress[1] | 213 | 55 | 5.39 e-14 | 4.01 | (Voskuil et al., 2011) |
| Nitric Oxide stress[1] | 237 | 48 | 2.13 e-08 | 2.83 | (Voskuil et al., 2011) |

[1]Considered significant at $p < 0.05$ and odds ratio > 1. Significantly overlapped pairs are highlighted in grey.

[2]$\log_2$ fold change> 1 or <-1 and $p$ value< 0.01 is considered differentially expressed genes in stress conditions.

**Table S5. Susceptibility of *M. tuberculosis* strains to diverse anti-TB drugs under *in vitro* aerobic growth conditions.**

| | $MIC_{90}$ (µg/ml)[1] | |
|---|---|---|
| Anti-TB drugs | *H37Rv* | *H37Rv-LbNox* |
| Isoniazid | 0.06 | 0.03 |
| Rifampicin | 0.008 | 0.008-0.016 |
| Ethambutol | 2.5-5 | 5 |
| Bedaquiline | 0.5 | 0.25-0.5 |

[1]Data shown are the result of two independent experiments performed in quadruplicate.

**Table S6. Mutant prevention concentration (MPC) of moxifloxacin with NAC against MDR *M. tuberculosis* strain NHN1664.**

| Moxifloxacin concentration μM (fold MIC[a]) | NAC Concentration (mM) | Number of Resistant colonies on drug containing 7H11[b] | | | | Mutation Frequency with Moxifloxacin |
|---|---|---|---|---|---|---|
| 1 (2X) | 0 | 406 | 405 | | | $1.6\times10^{-7}$ |
| | 1 | 45 | 55 | 70 | 124 | $3.0\times10^{-8}$ |
| | 2 | 16 | 7 | 4 | 1 | $2.8\times10^{-9}$ |
| 2 (4X) | 0 | 4 | 6 | 17 | 39 | $6.7\times10^{-9}$ |
| | 1 | 0 | 0 | 0 | 5 | $<4.0\times10^{-10}$ - $2.0\times10^{-9}$ |
| | 2 | 0 | | | | $<4.0\times10^{-10}$ |
| 4 (8X[c]) | 0 | 0 | | | | $<4.0\times10^{-10}$ |
| | 1 | 0 | | | | $<4.0\times10^{-10}$ |
| | 2 | 0 | | | | $<4.0\times10^{-10}$ |

[a]fold MIC with respect to MIC of H37Rv (0.5 μM)
[b]Number of input bacteria = $2.5\times10^{9}$ per plate. Data are from two independent experiments performed in duplicate.
[c]MPC for drug alone.

**Table S7. N-acetyl cysteine decreases recovery of *M. tuberculosis* resistant mutants in infected BALB/c mice.**

| | UT[a] | NAC[b] | MOXI[c] | NAC+MOXI |
|---|---|---|---|---|
| Total Mice/ group | 6 | 6 | 6 | 6 |
| Number of mice harboring resistant colonies | 1 (17%) | 2 (33%) | 4 (67%) | 1 (17%) |
| Total Resistant colonies/ group | 1 | 2 | 8 | 1 |
| Average resistant colonies/ mouse[d] | 0.17 | 0.33 | 1.33 | 0.17 |

[a]6 mice were administered with water (vehicle control).
[b]500 mg/ kg body weight NAC was given once daily for a period of 2 weeks by oral gavage.
[c]50 mg/ kg body weight moxifloxacin (MOXI) was given once daily for a period of 2 weeks by oral gavage.
[d]average resistant colonies/ mouse = total number of resistant colonies in a treatment group divided by total number of mice in that group.

#### Supplementary Figures

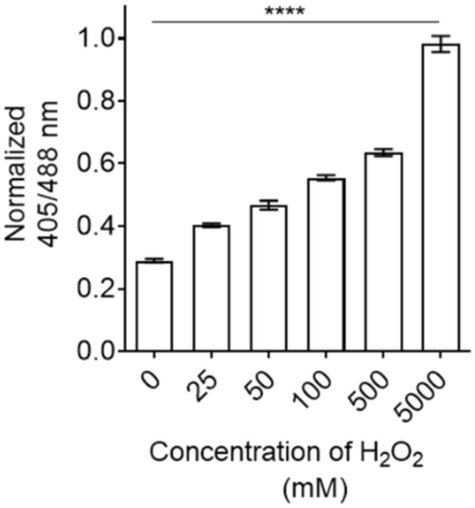


**Fig. S1. Mrx1-roGFP2 measures redox changes in *M. tuberculosis* in response to $H_2O_2$.** *M. tuberculosis* expressing Mrx1-roGFP2 was treated with the indicated concentrations of $H_2O_2$ for 5 min, and the ratiometric sensor response was measured by flow cytometry. A 2-fold increase in biosensor oxidation inside cells corresponds to biosensor oxidation in cells treated with 500 µM of $H_2O_2$ when compared to untreated cells. Error bars represent standard deviation from the mean. Data represent at least two independent experiments performed in at least duplicate. Statistical significance was analyzed over untreated control by one-way ANOVA analysis (****$p < 0.0001$).

Since $H_2O_2$ alone (0.5–5 mM) does not directly oxidize the Mrx1-roGFP2 (Bhaskar et al., 2014), the data suggest that $H_2O_2$-mediated oxidation of MSH to MSSM occurs via an intermediary enzyme (mycothiol peroxidase or peroxiredoxins), resulting in biosensor oxidation. These results, along with previous findings with Mrx1-roGFP2, show a direct relationship between changes in biosensor ratio and mycobacterial redox physiology (Bhaskar et al., 2014; Mishra et al., 2019; Mishra et al., 2017; Nambi et al., 2015).

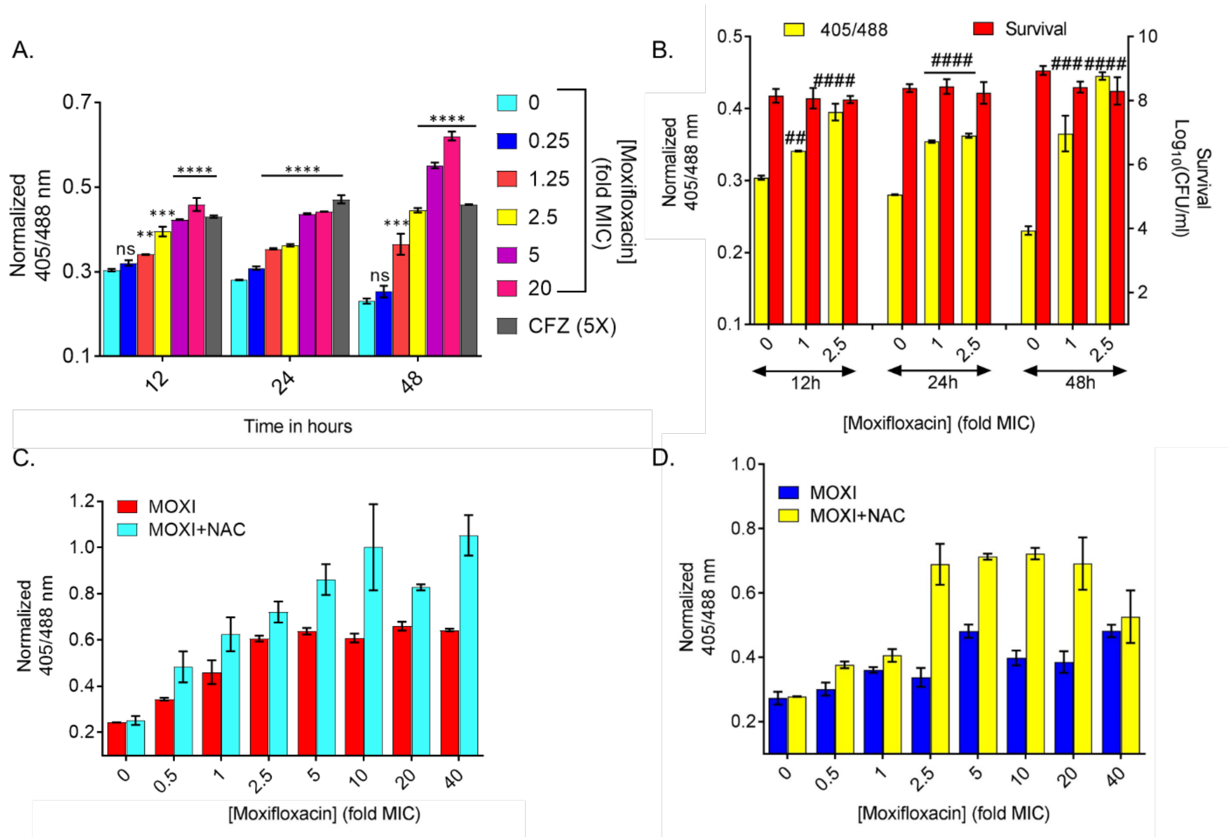


**Fig. S2. Effect of moxifloxacin and NAC on redox biosensor oxidation in *M. tuberculosis* cultures.** (**A**) Effect of moxifloxacin on redox biosensor. Log-phase *M. tuberculosis*-roGFP2 cultures were treated with the indicated concentrations of moxifloxacin (MOXI; 1X MIC= 0.5 µm), and the ratiometric response of the biosensor was determined at the indicated times. Moxifloxacin at sub-inhibitory concentrations did not induce oxidative stress. Clofazimine (CFZ at 5X MIC = 2.5 µg/mL) served as a positive control. (**B**) Time-resolved kinetics of ROS production and survival of *M. tuberculosis* during an early phase of moxifloxacin treatment. Difference in survival (red bars) is not significant at 12 h and 24 h. (**C**) Effect of NAC on biosensor signal. *M. tuberculosis*-roGFP2 was grown to $OD_{600}$ = 0.2-0.3, pretreated with 1 mM NAC for 1 h, and then treated with the indicated concentrations of moxifloxacin for 48 h. (**D**) Effect of NAC on *M. tuberculosis* NHN1664 expressing the biosensor. Conditions were as in panel C.

Data in this figure show that moxifloxacin treatment oxidizes, and increases, the redox biosensor signal and that NAC further increases the biosensor signal. The NAC-only control is shown as zero moxifloxacin. Data represent two independent experiments performed in duplicate. Error bars represent standard deviation from the mean. *p* was determined by unpaired two-tailed student's t-test analyzed relative to an untreated control. (* $p$ <0.05, ** $p \le 0.01$ *and* *** $p \le 0.001$ relative to the untreated control; ns indicates not significant). (## $p \le 0.01$, ### $p \le 0.001$ and #### $p < 0.0001$)

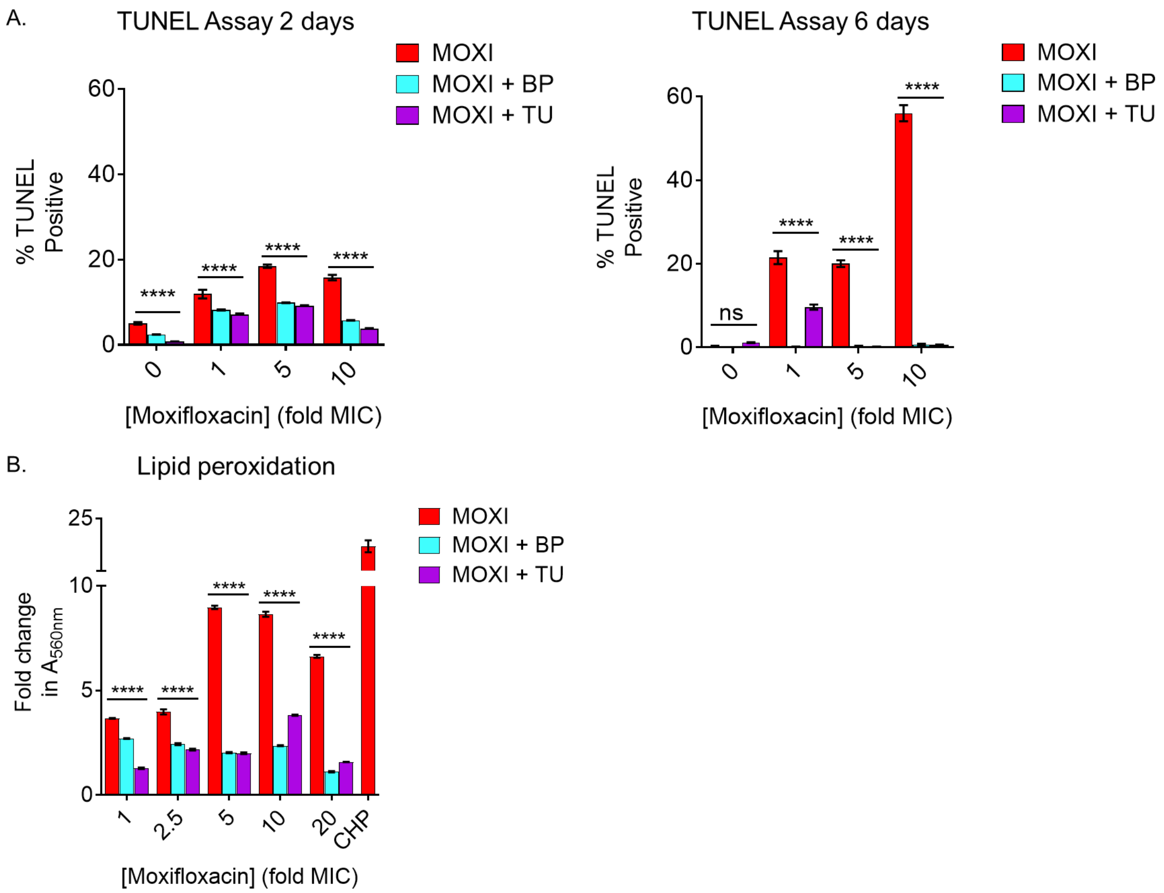


**Fig. S3. Moxifloxacin treatment increases markers of oxidative stress (DNA double-strand breaks and lipid hydroperoxides) in *M. tuberculosis*.** Exponentially growing *M. tuberculosis* cultures were either left untreated or treated with 250 μM bipyridyl (BP) for 15 min or with 10 mM thiourea (TU) for 1 h prior to addition of the indicated concentrations of moxifloxacin (MOXI) (1X MIC= 0.5 μM) for either 2 days or 6 days. **(A)** TUNEL Assay for measuring DNA double-stranded breaks using flow cytometry. **(B)** Lipid hydroperoxides were isolated after 2 days of MOXI treatment and measured after incubating with FOX2 reagent for 6 h. Data are normalized to culture cell density ($OD_{600}$) and represented as fold change in absorbance at 560 nm over respective untreated controls. Error bars represent standard deviation from the mean. Data represent at least two independent experiments performed in at least duplicate. Statistical significance was calculated with drug-alone group and with the bipyridyl/thiourea-only treatment groups with the drug + bipyridyl/thiourea groups by two-way ANOVA analysis; (****$p$ < 0.0001, ns indicates not significant).

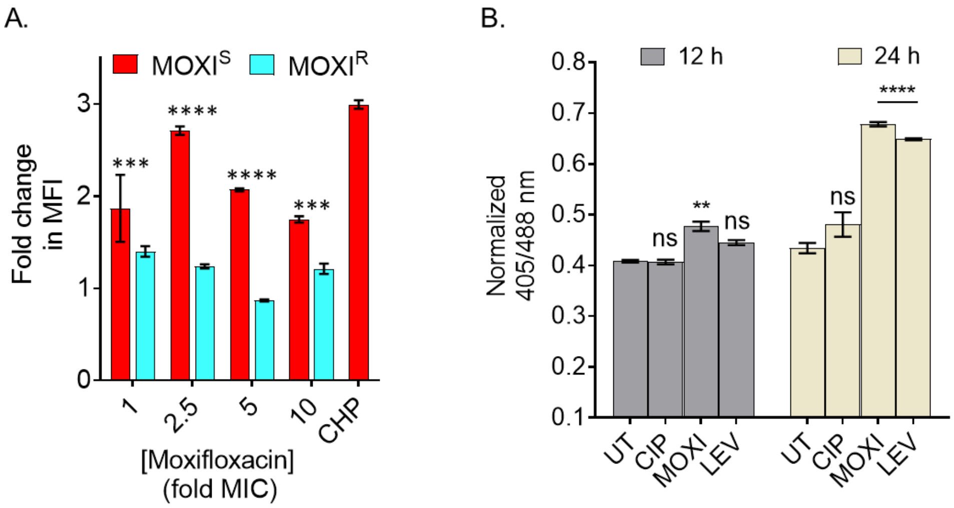


**Fig. S4. Effect of various fluoroquinolones on ROS production in *M. tuberculosis*. (A)** Strain *M. tuberculosis*-roGFP2 (moxifloxacin sensitive (MOXI$^S$); 1X MIC = 0.5 µM) and a moxifloxacin-resistant *M. tuberculosis* strain (MOXI$^R$); 1X MIC = 1.25 µM) were exposed to the indicated concentrations of moxifloxacin (1X MIC = 0.5 µM) for 48 h, and ROS were quantified by flow-cytometry using CellROX Deep Red dye. An oxidant, cumene hydroperoxide (CHP; 10 mM), served as a positive control. Data represent fold change in median fluorescence intensity of the dye compared to an untreated control. **(B)** Exponentially growing log-phase *M. tuberculosis*-roGFP2 cultures were treated with 2.5X MBC of moxifloxacin (MOXI), ciprofloxacin (CIP), or levofloxacin (LEV), and the ratiometric response of the biosensor was determined at 12 h and 24 h of treatment. Ciprofloxacin, which has a higher MBC, does not induce oxidative stress. Error bars represent standard deviation from the mean. Data shown are representative of two independent experiments performed in duplicate. Statistical considerations were as in Fig. S2.

These data address the robustness of the relationship between ROS and survival of *M. tuberculosis* following fluoroquinolone treatment. For various fluoroquinolones, killing is known to be greater for those with a lower MIC; thus, the ROS data fit with killing. For the resistant mutant, prevention of primary-lesion formation blocks the downstream death process; a lower ROS signal is the expected result.

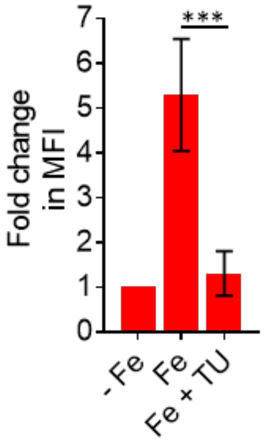


**Fig. S5. Increased free iron (Fe) concentration leads to ROS generation in *M. tuberculosis*.** Fe-depleted bacterial cultures (-Fe) were supplemented with 80 µM ferric chloride ($FeCl_3$) in the presence or absence of an ROS scavenger, thiourea (TU; 10 mM), for 4 days, and ROS were quantified by flow-cytometry using CellROX Deep Red dye. Data represent fold change in median fluorescence intensity (MFI) of the dye over untreated control (-Fe). Error bars represent standard deviation of the mean. Data shown are representative of two independent experiments performed in duplicate. Statistical considerations were as in Fig. S2.

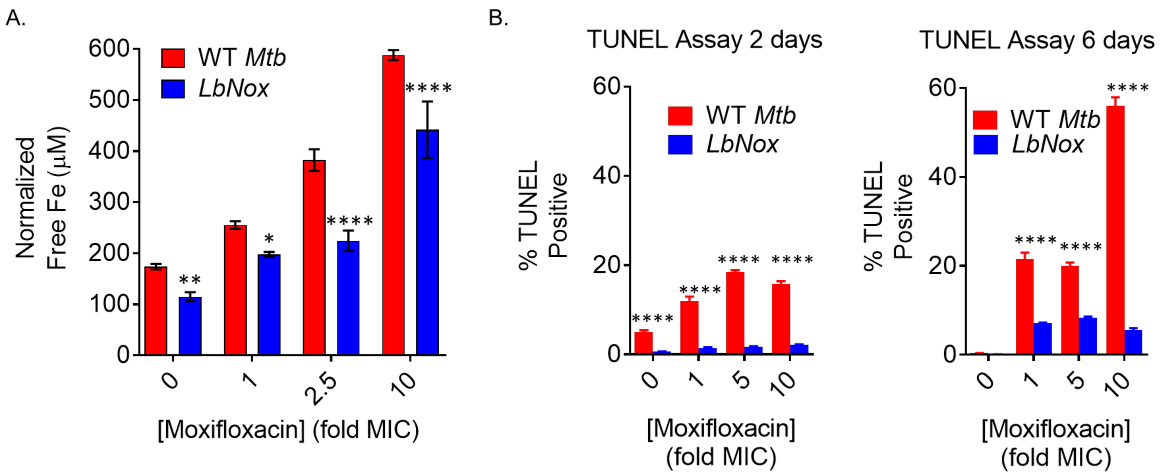


**Fig. S6. Moxifloxacin treatment increases free Fe pools and DNA breaks in *M. tuberculosis* that are decreased by NADH dissipation. (A)** Detection of free Fe levels. Exponentially growing *M. tuberculosis* cultures were treated with the indicated concentrations of moxifloxacin (1X MIC=0.5 µM) for 2 days, and free Fe levels were determined by a ferrozine-based colorimetric assay. **(B)** TUNEL Assay for measuring DNA double-stranded breaks using flow cytometry. Results were compared with drug-treated, wild-type *M. tuberculosis*. Error bars represent standard deviation from the mean. Data shown are representative of two independent experiments performed in triplicate. Statistical considerations were as in Fig. S2.

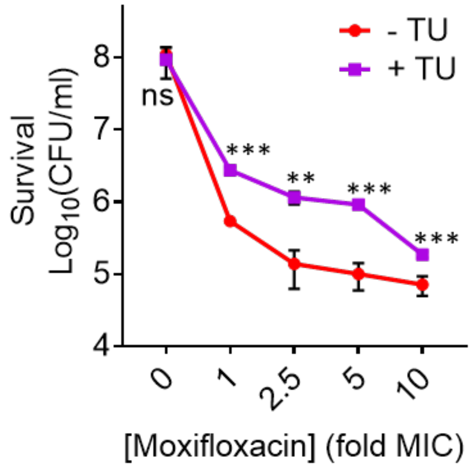


**Fig. S7: Reduction in moxifloxacin lethal activity by the ROS scavenger thiourea.** Exponentially growing *M. tuberculosis* cells were either left untreated or treated with 10 mM TU for 1 h prior to addition of the indicated concentrations of moxifloxacin (1X MIC=0.5 µM) for 2 days followed by determination of CFU on drug-free 7H11 plates. Error bars represent standard deviation from the mean. Data shown are representative of two independent experiments performed in duplicate. Statistical considerations were as in Fig. S2.

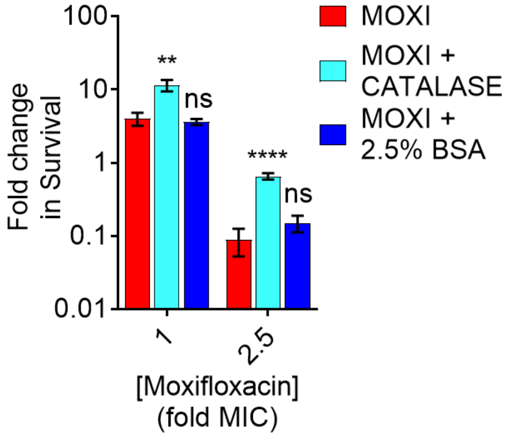


**Fig. S8. Addition of extra bovine serum albumin to agar plates does not protect *M. tuberculosis* from moxifloxacin-mediated lethality.** *M. tuberculosis* cultures were treated with 1X and 2.5X MIC of moxifloxacin (MOXI; 1X MIC= 0.5 µM) for 48 h, washed, and then plated on 7H11 agar without catalase, with catalase (17.5 U/mL of 7H11 agar), or with 2.5% bovine serum albumin (BSA; 5-fold higher than normal BSA content in 7H11 agar). Fold change in survival was calculated relative to untreated control. Error bars represent standard deviation from the mean. Statistical significance was calculated between drug-alone group with drug + catalase or drug + 2.5% BSA groups. Statistical considerations were as in Fig. S2.

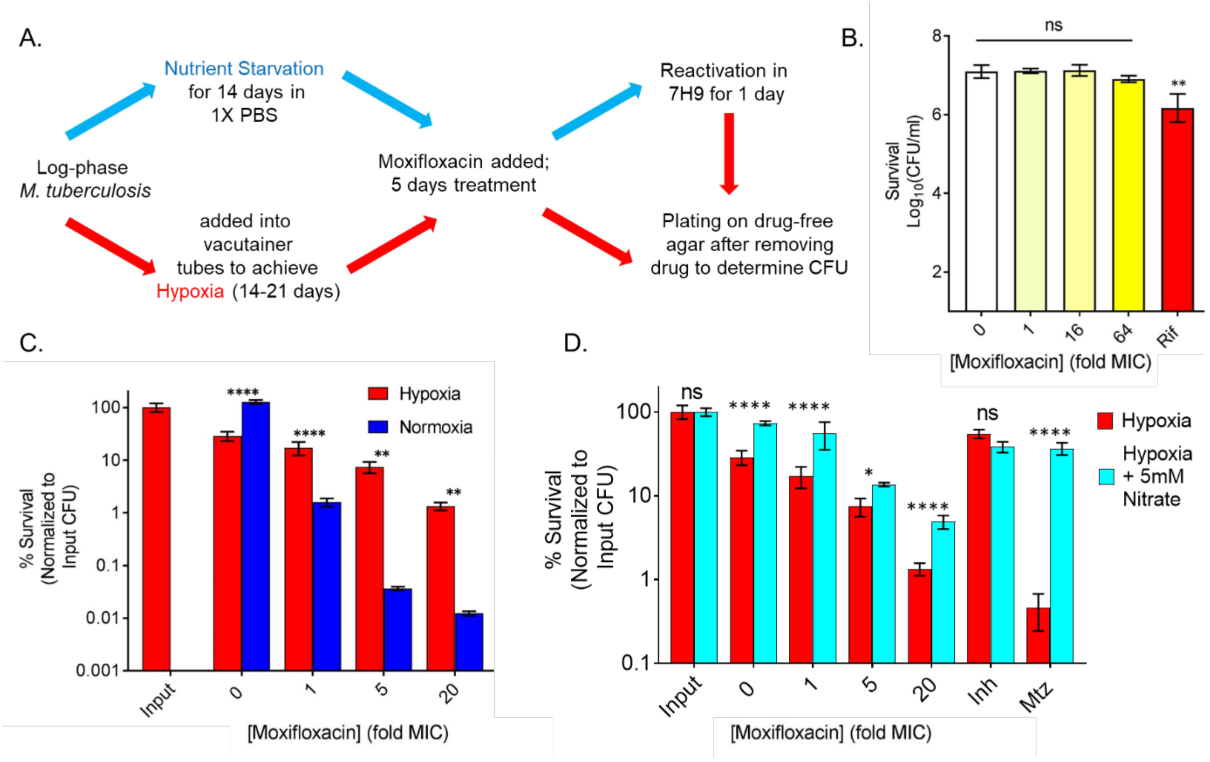


**Fig. S9. Moxifloxacin lethality during nutrient starvation and hypoxia.** (**A**) Experimental plan for examining effects of moxifloxacin under nutrient starvation (red arrows) and hypoxia (green arrows). (**B**) Effect of nutrient starvation. *M. tuberculosis* cultures were starved for nutrients for 14 days and then treated with moxifloxacin (1X MIC= 0.5 μM) for 5 days before determination of survival. Rifampicin (Rif; 25 μM) served as a drug control. (**C**) Effect of hypoxia. Survival of *M. tuberculosis* under hypoxia and aerobic culture conditions (normoxia) when treated with the indicated concentrations of moxifloxacin for 5 days, followed by CFU determination. (**D**) Effect of nitrate during hypoxia. Sodium nitrate (5 mM) was added when cultures were placed in Vacutainer tubes; treatment conditions were as indicated in Materials and Methods. Survival of *M. tuberculosis* was measured as CFU after treatment with the indicated concentrations of moxifloxacin for 5 days. Isoniazid (Inh; 10 μM) and metronidazole (Mtz; 10 mM) were controls. Statistical considerations were as in Fig. S2.

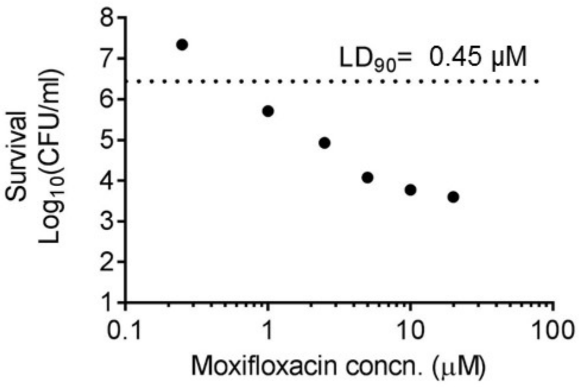


**Fig. S10. Determination of lethal dose ($LD_{90}$) of moxifloxacin with aerobically cultured *M. tuberculosis*.** Exponentially growing cultures of *M. tuberculosis* were treated for 5 days with the indicated concentrations of moxifloxacin after which aliquots were diluted and plated on drug-free agar and incubated for 3-4 weeks. The experiment was performed twice in triplicate. Dotted line indicates 90% reduction in CFU compared to input control (CFU at the time of drug addition); the corresponding concentration of moxifloxacin was taken as $LD_{90}$.

A

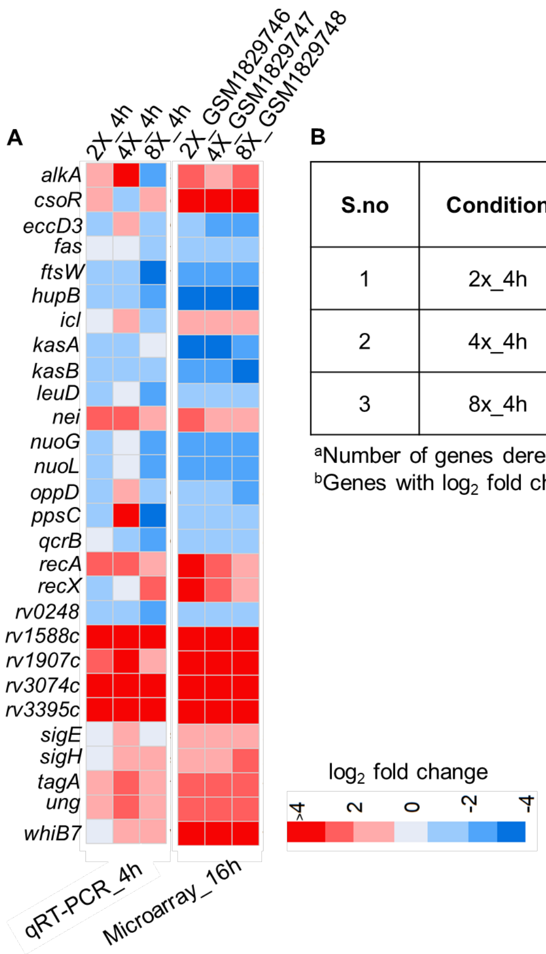


B

| S.no | Condition | Number of total genes in the condition | Overlap[a] | *p* value |
|---|---|---|---|---|
| 1 | 2x_4h | 28 | 18[b] | 1.30 e-16 |
| 2 | 4x_4h | 28 | 15[b] | 7.22 e-12 |
| 3 | 8x_4h | 28 | 21[b] | 2.47 e-22 |

[a]Number of genes deregulated at both time points.
[b]Genes with $\log_2$ fold change > 1 at 4 h is considered to be deregulated.

**Fig. S11. Heat map showing qRT-PCR validation of 28 randomly selected genes differentially regulated in the microarray data. A.** (qRT-PCR_4 h) Heat map depicting expression of genes ($\log_2$ fold change) for 4-h treatment with 2X, 4X, or 8X moxifloxacin (1X MIC = 0.4 μM) compared to untreated control using qRT-PCR analysis. (Microarray 16 h) indicates heat map depicting expression of these 28 genes ($\log_2$-fold change) for 16 h of 2X, 4X, or 8X moxifloxacin-treated *M. tuberculosis* compared to untreated control by microarray data. Color code for the fold change is shown (red: upregulated genes; blue: down-regulated genes). The expression pattern of the 28 differentially expressed genes at 4 h is similar to that of the 16-h treatment. **B.** Table showing *p* value for overlap analysis between genes deregulated at 4 h and 16 h after moxifloxacin treatment.

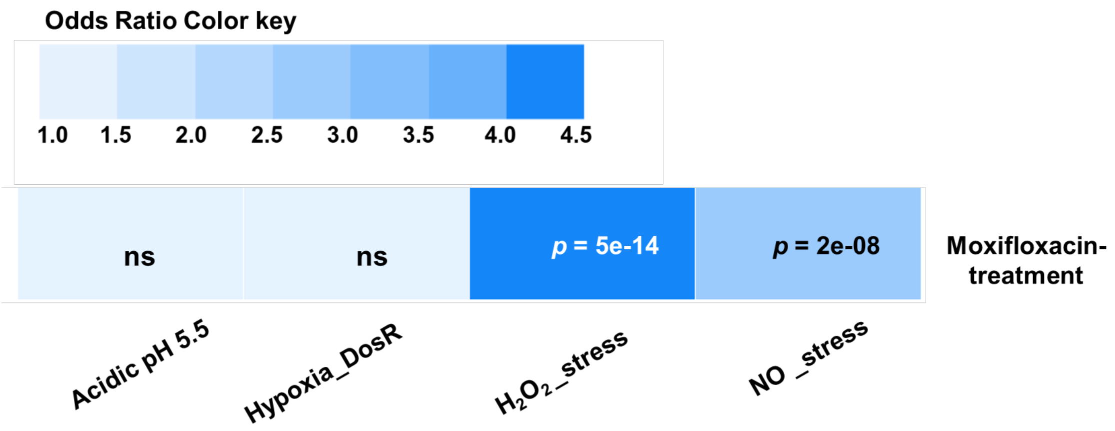


**Fig. S12. Heatmap showing differential gene overlap between moxifloxacin-exposed *M. tuberculosis* and various stress conditions.** Each column represents the overlap between the moxifloxacin-treatment and the individual stress condition. The *p*-value of the overlap is indicated for each pair. The color represents the odds ratio for the overlap. $H_2O_2$ stress showed significant overlap ($p$-value < 0.05) with a strong association (odds ratio > 1) with moxifloxacin stress. The acidic pH 5.5 and DosR regulated-genes in the hypoxic condition showed no significant overlap. (ns indicates not-significant).

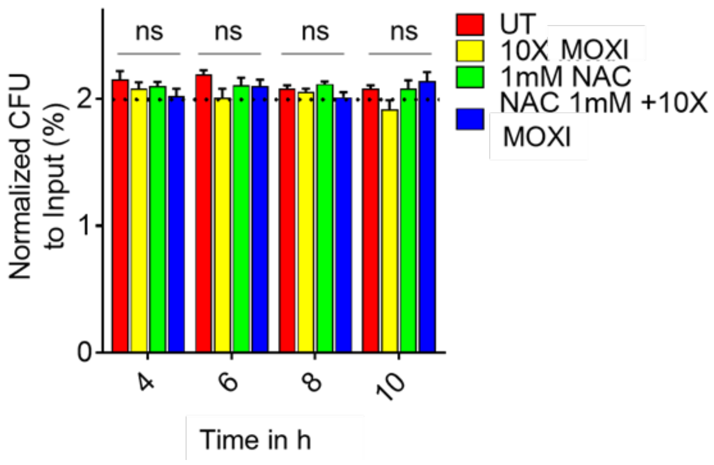


**Fig. S13. Decelerated respiration in *M. tuberculosis* upon moxifloxacin treatment is not due to bacterial death.** Bacterial cultures were prepared as for measurement of bioenergetics. They were then treated with moxifloxacin (MOXI; 10X MIC=5 μM) alone, NAC alone (1 mM), or the combination for 10 h. Samples were taken at the indicated times and plated for determination of CFU, which was normalized to CFU at the beginning of the experiment. Error bars represent standard deviation from the mean; ns indicates not significant. The data show that during the 10-h incubation, moxifloxacin and NAC had no significant effect on *M. tuberculosis* viability. Statistical considerations were as in Fig. S2.

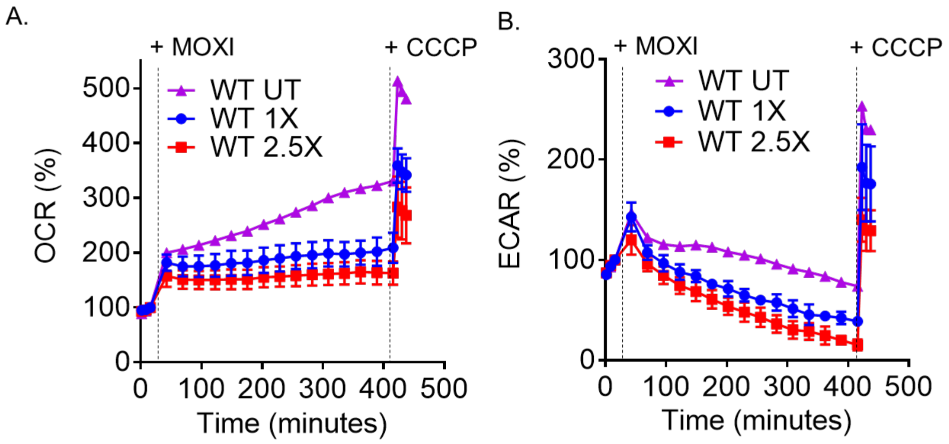


**Fig. S14. Low moxifloxacin concentrations decelerate respiration rate in *M. tuberculosis.*** (**A**) OCR (pmol/min), an indicator of oxygen consumption rate. Exponentially growing *M. tuberculosis* cultures were either left untreated (UT) or treated with 1X or 2.5X MIC of moxifloxacin (MOXI; 1X MIC=0.5 μM) for the indicated times; black dotted lines indicate the time when MOXI or NAC (1 mM) or CCCP (10 μM) were added. Determination was via Seahorse XFp Analyzer (**B**) ECAR (mpH/min), an indicator of $H^+$ production or extracellular acidification due to glycolytic and TCA flux. Determination was as in **A**. OCR and ECAR data represent percentage of third baseline value. Error bars represent standard deviation from the mean. Data shown represent two independent experiments performed in triplicate.

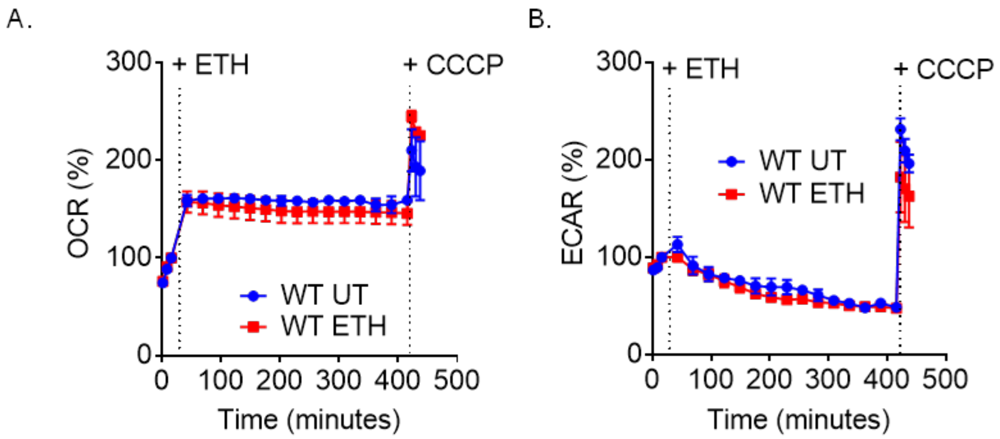


**Fig. S15. Ethambutol treatment does not affect respiration rate in *M. tuberculosis*.** (**A**) OCR (pmol/min), an indicator of oxygen consumption rate. Exponentially growing *M. tuberculosis* cultures were either left untreated (UT) or treated with 10X MIC of ethambutol (ETH; 1X MIC= 1 μg/ mL); black dotted lines indicate the time when ETH or CCCP (10 μM) were added to cells. Determination was via Seahorse XFp Analyzer (**B**) ECAR (mpH/min), an indicator of $H^+$ production or extracellular acidification due to glycolytic and TCA flux. Determination was as in *A*. OCR and ECAR data represent percentage of third baseline value. Error bars represent standard deviation from the mean. Data shown are representative of two independent experiments performed in triplicate.

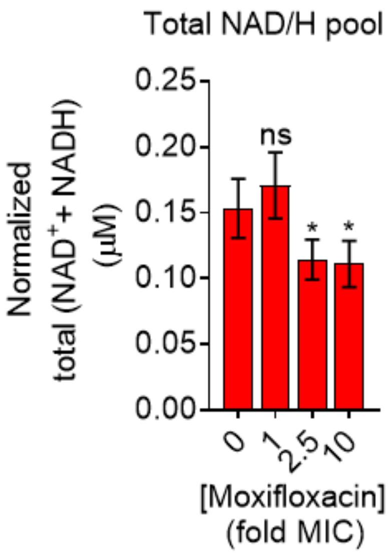


**Fig. S16. Moxifloxacin decreases total NAD/H (NADH+NAD$^+$) in *M. tuberculosis*.** Detection of total NADH + NAD$^+$ levels. Exponentially growing *M. tuberculosis* was treated with the indicated concentrations of moxifloxacin (1X MIC=0.5 µM) for 2 days, and total NADH+ NAD$^+$ levels were determined by an alcohol dehydrogenase-based redox cycling assay. Error bars represent standard deviation from the mean. Data represent at least two independent experiments performed in at least duplicate. *p* was determined by unpaired two-tailed student's t-test analyzed relative to untreated control. (*; $p < 0.05$, ns indicates not significant).

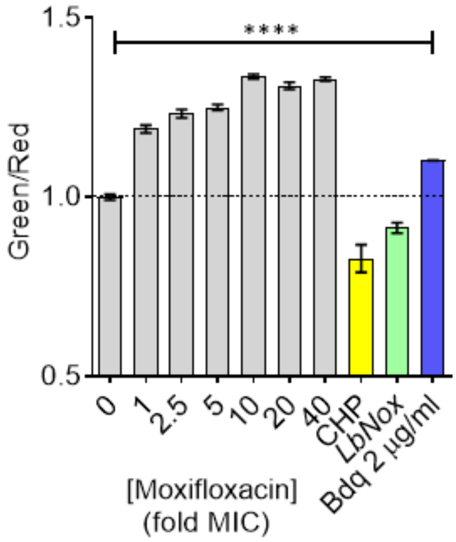


**Fig. S17. Moxifloxacin increases NADH/NAD⁺ ratios in *M. tuberculosis* detected by Peredox.** *M. tuberculosis* H37Rv, expressing the Peredox biosensor, was treated with indicated concentrations of moxifloxacin (1X MIC= 0.5 μM) for 48 h; green (Ex/Em. = 405/510 nm) and red fluorescence (Ex/Em. = 560/615 nm) were measured by FACS. The green/red ratio corresponds to NADH/NAD+ levels in the bacterial culture. NADH-responsive Rex protein is fused to T-sapphire which fluoresces based on dynamic NADH:NAD+ levels; mCherry fluorescence is a normalizing control. Untreated *LbNox* strain, cumene hydroperoxide (CHP; 1 mM), and bedaquiline (Bdq; 2 μg/ml)-treated cells expressing Peredox are controls. Error bars represent standard deviation from the mean. Data are representative of two independent experiments performed in duplicate. Statistical significance was determined by one-way ANOVA followed by Dunnett's test; ****$p < 0.001$.

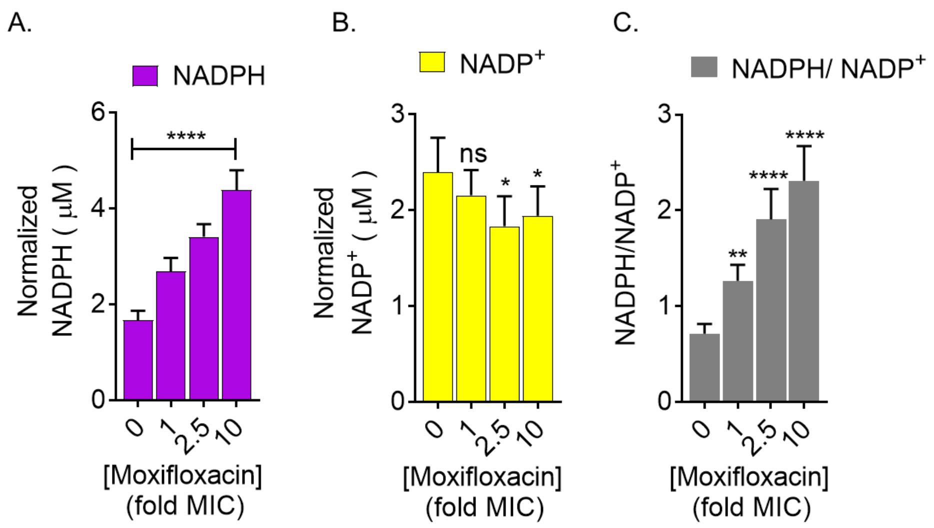


**Fig. S18. NADP+ and NADPH levels in moxifloxacin-treated *M. tuberculosis.*** Detection of **(A)** NADPH, **(B)** $NADP^+$, and **(C)** NADPH/$NADP^+$ ratio in exponentially growing *M. tuberculosis* treated with the indicated concentrations of moxifloxacin (1X MIC=0.5 µM) for 2 days. NADPH and $NADP^+$ levels were determined by a glucose-6-phosphate dehydrogenase-based redox cycling assay. *p* was determined by unpaired two-tailed student's t-test analyzed relative to an untreated control. (* $p$ <0.05, ** $p \leq$ 0.01, **** $p \leq$ 0.0001, and ns indicates not significant).

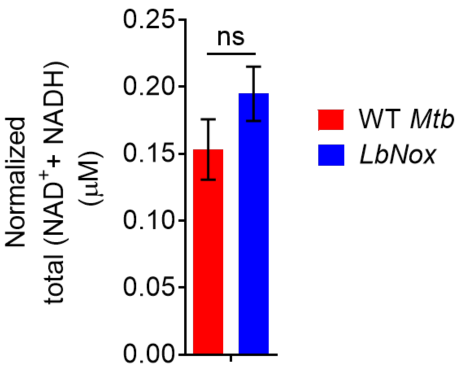


**Fig. S19. Total NAD/H (NADH+$NAD^+$) was not affected by overexpression of *LbNox* in *M. tuberculosis*.** *M. tuberculosis* cultures were grown to log-phase ($OD_{600}$ = 0.6-0.8), and total NADH+$NAD^+$ levels were determined by an alcohol dehydrogenase-based redox cycling assay. Error bars represent standard deviation of the mean. Data represent two independent experiments performed in at least duplicate. *p* was determined by unpaired two-tailed student's t-test analyzed relative to an untreated control (ns indicates not significant).

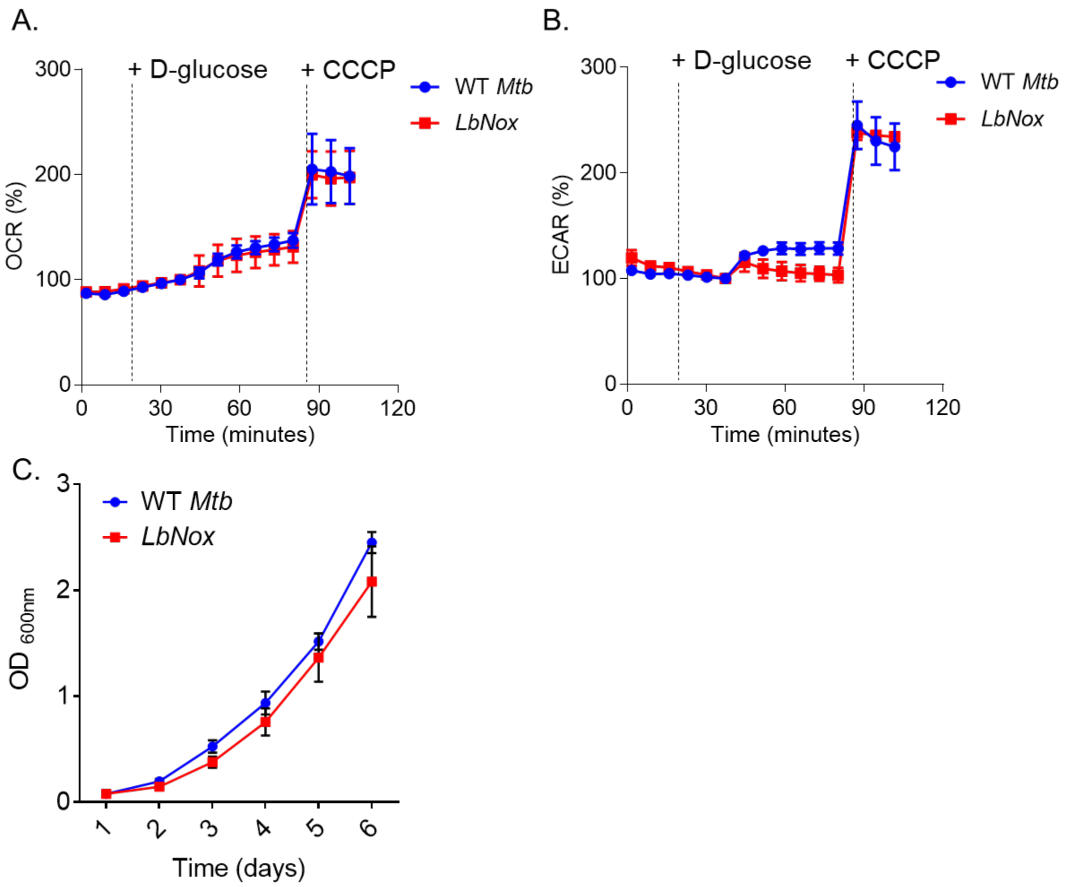


**Fig. S20. Constitutive expression of *LbNox* does not impair metabolism and growth rate of *M. tuberculosis*.** Exponentially growing wild-type *M. tuberculosis (*WT *Mtb)* and an *LbNox*-overexpressing strain were starved overnight for glucose. Changes in **(A)** Oxygen Consumption Rate (OCR) and **(B)** Extracellular Acidification Rate (ECAR) were quantified after addition of 2 mg/mL D-glucose and subsequently by 10 μM of the uncoupler CCCP using a Seahorse XF flux analyzer. Black dotted lines indicate the time when D-glucose or CCCP were added to cultures Data shown are representative of two independent experiments performed in triplicate. Error bars represent standard deviation from the mean. **(C)** Culture turbidity ($OD_{600}$) for wild-type (WT *Mtb*) *M. tuberculosis* and the *LbNox* strain in 7H9+ADS broth was measured. Data shown are the result of two independent experiments. Error bars represent standard deviation from the mean.

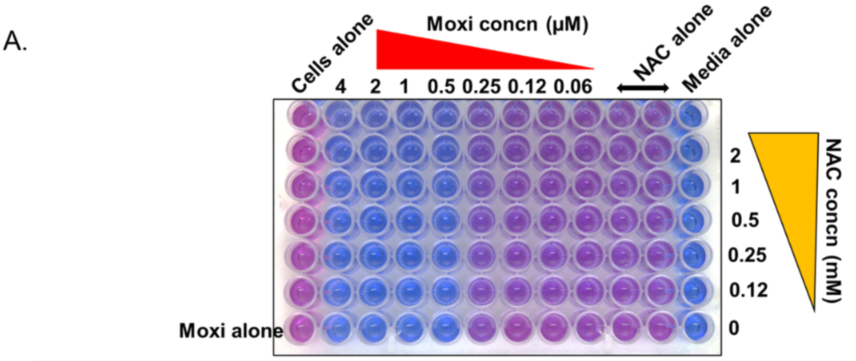


| | | | | | | | | | | |
|---|---|---|---|---|---|---|---|---|---|---|
| | | | | | | | | | only NAC | NAC concn (mM) |
| % inhibition | 110.8 | 109.5 | 108.7 | 104.9 | 55.8 | 51.4 | 38.6 | 40.5 | 40.7 | 2.00 |
| | 103.7 | 102.0 | 101.4 | 97.0 | 45.8 | 41.0 | 29.0 | 29.1 | 29.3 | 1.00 |
| | 101.8 | 100.5 | 99.8 | 94.8 | 37.9 | 33.1 | 22.4 | 23.9 | 29.7 | 0.50 |
| | 101.6 | 100.5 | 99.8 | 94.4 | 38.4 | 33.9 | 35.2 | 34.9 | 34.9 | 0.25 |
| | 101.8 | 100.6 | 100.0 | 95.1 | 39.1 | 31.9 | 22.2 | 34.5 | 34.4 | 0.13 |
| only moxi | 98.6 | 98.0 | 97.1 | 93.1 | 24.0 | 10.1 | 12.1 | 9.2 | | 0.00 |
| moxi concn (µM) | 4.00 | 2.00 | 1.00 | 0.50 | 0.25 | 0.13 | 0.06 | 0.03 | 0.00 | |

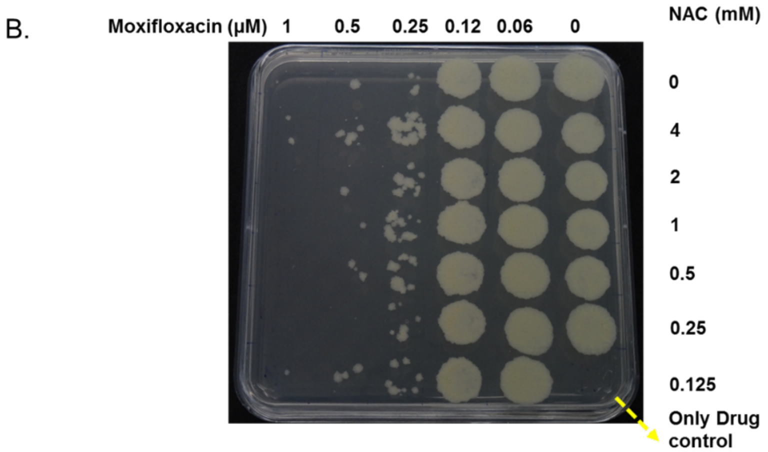


**Fig. S21. NAC has no effect on moxifloxacin MIC with *M. tuberculosis*. (A)** REMA assay**.** Color change from non-fluorescent blue resazurin to fluorescent pink resorufin by cellular metabolic activity indicates reduction of resazurin in the assay. % Inhibition of color change with respect to cells-only control is shown in the table. % Inhibition by moxifloxacin alone decreases with decreasing concentration, as shown in red. % Inhibition by NAC alone is shown in green. 90% inhibition is considered as MIC (= 0.5 µM for moxifloxacin alone; yellow highlight). No change in MIC is observed in the presence of NAC. **(B)** 7H11 Agar plate assay for MIC. 20 µL cells from MIC plate were regrown on 7H11 agar. NAC also had no effect on moxifloxacin MIC when measured by growth on drug-containing agar plates. Data shown are representative of two independent experiments.

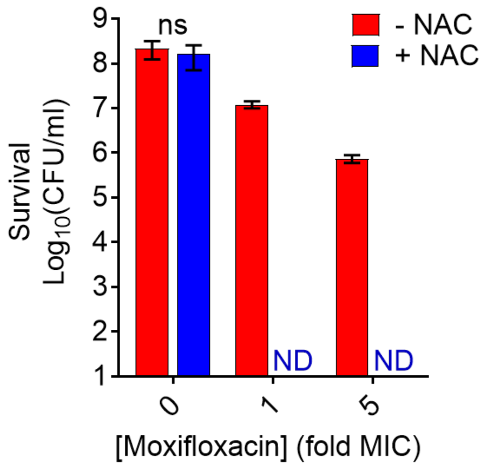


**Fig. S22. Addition of NAC after removal of moxifloxacin increases killing of *M. tuberculosis*.** In this assay for post-stressor death, *M. tuberculosis* cultures were treated with moxifloxacin (1X MIC= 0.5 µM) for 48 h, the drug was removed by washing, and cells were plated on drug-free 7H11 agar with or without 1 mM NAC. After incubation for 2 weeks, CFU were determined visually. ND indicates no colony was detected. Error bars represent standard deviation from the mean. Statistical significance was calculated between drug-alone group with drug + NAC group. Data shown are the result of two independent experiments performed in duplicate. Statistical considerations were as in Fig. S2.

This experiment is the complement to the reduction in killing seen when catalase is present on agar to lower ROS in cells. The result strongly supports the idea that ROS, once levels pass a threshold, become self-sustaining: the presence of the initial stressor is no longer required. This is the result expected from a stress-mediated death pathway.

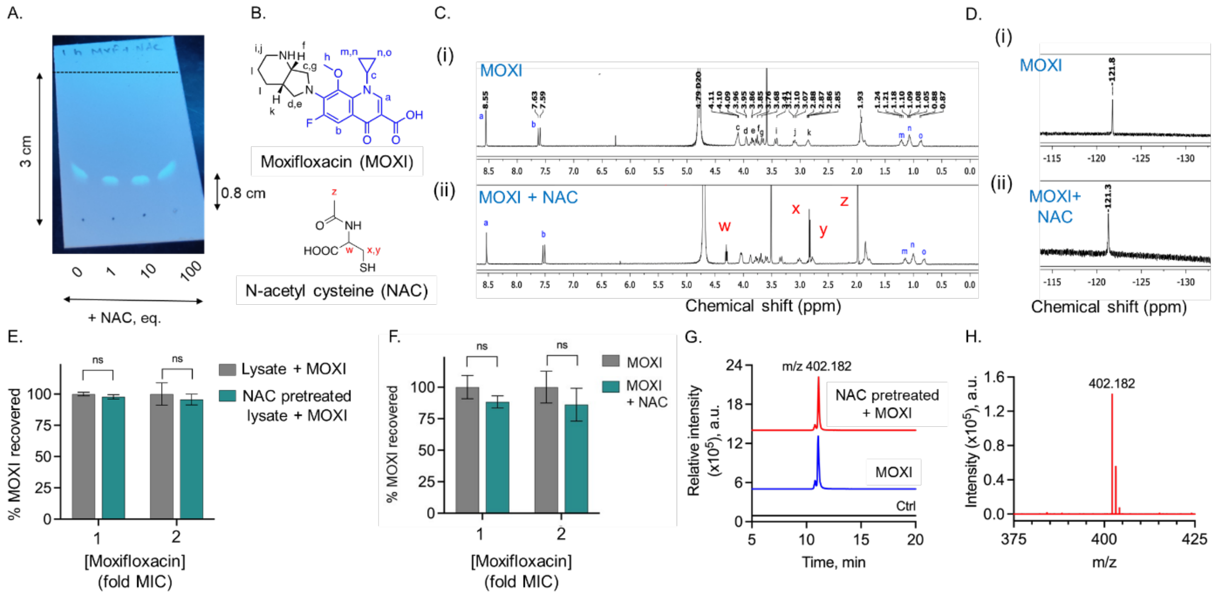


**Fig. S23. Several tests argue against formation of adducts between NAC and moxifloxacin (MOXI).**

**(A)** Thin layer chromatography (TLC) analysis of the reaction mixture containing MOXI ($R_f$ = 0.26) alone or with NAC in pH 7.4 buffer at 37 °C. The solvent system used was 1:9 methanol/ chloroform. and the TLC plate was visualized in a UV chamber at 254 nm. Reaction time was 60 min. The lanes indicate (1) authentic MOXI ($R_f$ = 0.26), (2) MOXI + NAC (1:1), (3) MOXI + NAC (1:10), and (4) MOXI + NAC (1:100). Although the result shows curvature, the position of the moxifloxacin signal is unchanged, and no new signal representing a putative adduct was seen.

**(B)** Structures of MOXI and NAC.

**(C, D)** NMR analysis. Stacked spectra of (C) $^{1}$H-NMR and (D) $^{19}$F-NMR acquired after 1 h incubation of (i) MOXI alone and (ii) MOXI with NAC (1:1) in deuterated phosphate buffer (pH = 7.4, 10 mM) at 37 °C. No new signal corresponding to putative MOXI-NAC adduct was observed. The singlet and doublet peaks corresponding to characteristic ‘a’ and ‘b’ protons (highlighted in blue) of MOXI at $\delta_H$ 8.55 and 7.61 ppm in the $^{1}$H-NMR spectrum remain largely unchanged in the presence of NAC. Similarly, no significant change in the $^{19}$F-NMR spectrum of MOXI was created by the presence of NAC.

**(E)** Monitoring the amount of MOXI recovered by a fluorescence-based assay. *M. smegmatis* lysates (1 mg/mL; protein concentration in lysate as determined by Bradford assay) were treated with MOXI alone or pre-treated with NAC (1 mM) followed by MOXI. Fluorescence measurements were carried out after 24 h of incubation at the indicated MIC concentrations (1X and 2X; 0.25 μM and 0.5 μM) for MOXI. Error bars represent standard deviation. Data shown are representative of two independent experiments, each performed in triplicate. (ns indicates not significant).

**(F)** Insignificant effect of NAC on recovery of moxifloxacin from *M. smegmatis*. Wild-type *M. smegmatis* cultures were pretreated with 1 mM NAC for 1 h and then moxifloxacin was added to either 1X or 2X MIC for a 48-h incubation. Detection of moxifloxacin was by fluorescence. Reduction of moxifloxacin recovery by NAC was insignificant. Error bars represent standard error of the mean. Data shown are representative of two independent experiments performed in triplicate. (ns indicates not significant).

**(G)** LC-MS based quantitative determination of MOXI in WT *M. smegmatis*. Extracted ion chromatograms (EIC) show no NAC-moxifloxacin adduct when obtained from treated *M. smegmatis*. A signal at m/z 402.1824 corresponded to MOXI (expected, $[M+H]^+$ = 402.1824; observed $[M+H]^+$ = 402.1813) recovered in lysates obtained from lysis of NAC (1 mM)-untreated and NAC-pretreated wild-type *M. smegmatis* challenged with 2x MIC (0.5 µM) of MOXI for 48 h. Ctrl refers to bacteria alone.

(**H**) LC-MS traces (MRM-HR) for MOXI (m/z = 402.182) recovered in NAC-treated wild-type *M. smegmatis* incubated with MOXI (2x MIC). Nearly quantitative recovery of MOXI (>85%) was observed under these conditions.

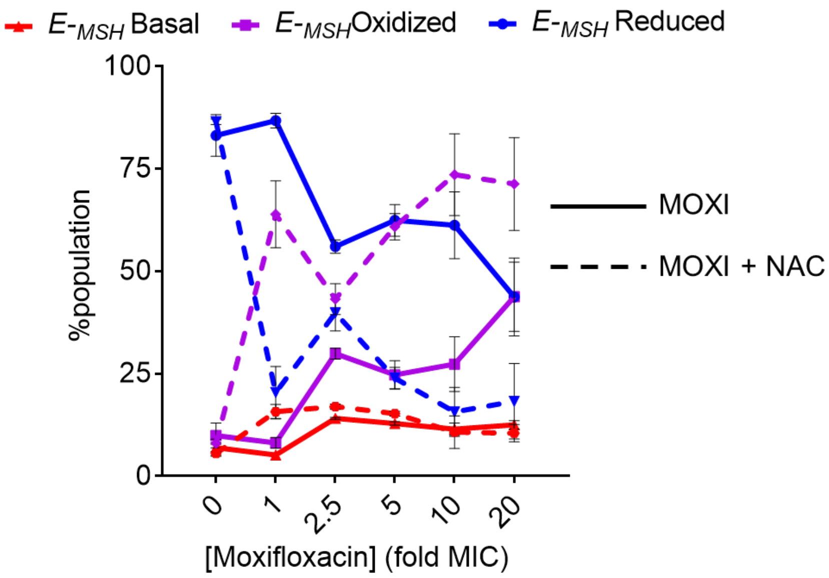


**Fig. S24. Moxifloxacin-induced oxidative shift in $E_{MSH}$ of *M. tuberculosis* NHN1664 during infection of THP-1 cells.** THP-1 macrophage-like cells, infected at an MOI of 1:10 with the MDR strain NHN1664 expressing Mrx1-roGFP2, were treated with the indicated concentrations of moxifloxacin (1X MIC= 0.5 µM) in the presence or absence of NAC (1 mM) immediately after infection. They were then incubated for 24 h. Approximately 10,000 infected macrophages were analyzed by flow cytometry to quantify changes in *M. tuberculosis* subpopulations displaying redox heterogeneity.

The data show that NAC induces an oxidative shift in an MDR strain that lowered the drug-tolerant $E_{MSH}$-reduced subpopulation (blue line in line graph), similar to that seen with infection with the drug-sensitive laboratory strain *M. tuberculosis* H37Rv.

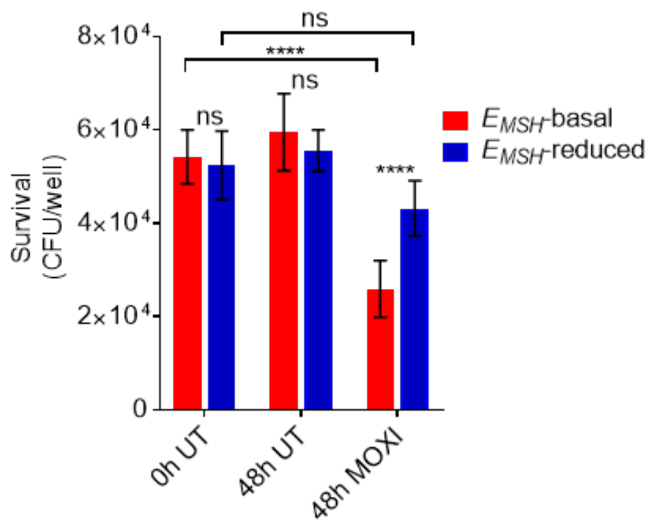


**Fig. S25. Intramacrophage *$E_{MSH}$*-reduced *M. tuberculosis* shows reduced killing by moxifloxacin lethality.** Infected macrophages were sorted by flow cytometry into those harboring *M. tuberculosis* at an $E_{MSH}$-basal and those with *M. tuberculosis* at an $E_{MSH}$-reduced redox state. The two groups were exposed to 3X MIC of moxifloxacin (MOXI; 1X MIC= 0.5 µM) or left untreated (UT) for 48 h. Macrophages were then lysed, and the bacillary load was determined by CFU enumeration by incubation of bacteria on agar plates. $E_{MSH}$ was determined as described in Supplementary Methods. Error bars represent standard deviation from the mean. Data shown are representative of two independent experiments performed in at least triplicate. *p* was determined by unpaired two-tailed student's t-test (****$p < 0.0001$, ns indicates not significant).

These data show that when the redox state of intracellular *M. tuberculosis* is reduced ($E_{MSH}$-reduced), moxifloxacin lethality is significantly decreased, while $E_{MSH}$-basal bacteria remain sensitive to the drug.
